# Natural language supervision with a large and diverse dataset builds better models of human high-level visual cortex

**DOI:** 10.1101/2022.09.27.508760

**Authors:** Aria Y. Wang, Kendrick Kay, Thomas Naselaris, Michael J. Tarr, Leila Wehbe

## Abstract

Advances in neural networks have been catalyzed by joint training on images and natural language, increased dataset sizes, and data diversity. We explored whether the same factors support similar improvements in predicting visual responses in the human brain. We used models pre-trained with Contrastive Language-Image Pre-training (CLIP) – which learns image embeddings that best match text embeddings of image captions from diverse, large-scale datasets – to study visual representations. We built voxelwise encoding models based on CLIP image features to predict brain responses to real-world images. ResNet50 with CLIP explained up to *R*^2^ = 79% of variance in individual voxel responses in held-out test data, a significant increase from models trained only with image/label pairs (ImageNet trained ResNet) or text (BERT). Comparisons across different model backbones ruled out network architecture as a factor in performance improvements. Comparisons across models that controlled for dataset size and data diversity demonstrated that language feedback along with data diversity in larger datasets are important factors in explaining neural responses in high-level visual brain regions. Visualizations of model embeddings and Principal Component Analysis (PCA) revealed that our models capture both global and fine-grained semantic dimensions represented within human visual cortex.

---

Advances in deep learning have sparked a revolution in machine intelligence^1^ and, in parallel, a revolution in how we explain brain recordings. Heretofore unaccounted for brain responses can now be predicted by deep neural networks that share task goals and learned representations with natural systems^2–4^. However, this endeavour has been limited by the fact that most models used for brain prediction learn a low dimensional task objective (e.g., categorization) and are based on pre-training with ImageNet^5^. In contrast, natural vision solves multiple tasks and has evolved over millions of years, incorporating diverse perceptual, conceptual, and language sources^6–9^. A major challenge for understanding biological systems is to consider such multimodal sources in network training, for example, by including incorporating complex datasets that capture human-relevant information.

We have recently seen dramatic performance improvements in both vision and language tasks for state-of-the-art models. These advances may be attributed, in part, to learning more complex human semantics from multiple modalities that help delineate what is important in training sets that are larger and more diverse than those used in earlier models^10–14^. Language is particularly effective in drawing attention to human semantics in training data in that language is generated by humans and has evolved to highlight aspects of the world that are behaviorally relevant^15^. At the same time, larger training set sizes provide more (and possibly better) examples for high-dimensional supervisory signals such as natural language^16^. Reinforcing the importance of these factors, we also see dramatic improvements in our ability to explain aspects of human vision using these same, state-of-the-art models.

We took models using “Contrastive Language-Image Pre-training” or “CLIP”^10^ as representative of the class of models that leverage supervision from natural language (image captions) for vision and from vision (scene images) for language^11–13,17^. CLIP models are trained with image/caption pairs, learning separate image and text encoders from scratch that encode each pair from the training data with similar representations at the final layer. As compared to previous multimodal models (e.g., VisualBERT^18^, LXMERT^19^), multimodal loss signals in the final layer are propagated through earlier layers of both the visual and language encoders. As compared to earlier models, model learning with CLIP may be more similar to human visual learning, where top-down knowledge influences the earliest layers of the visual pathway^20,21^. As such, CLIP is an attractive model for studying brain prediction along many dimensions. CLIP’s joint natural language and image pre-training and large, diverse training set better capture fine-grained human visual experience. Moreover, the versatile CLIP scheme allows us to explore the impact of different model architectures, while further controlled comparisons using related models and datasets likewise allow us to explore the impact of dataset size and diversity.

We extracted network representations from neural network models trained with CLIP, including ResNet*_CLIP_*and ViT*_CLIP_* and from several single modality models: ImageNet^5^ pre-trained ResNet50^22^ (“ResNet*_ImageNet_*”) and BERT^23^ (associated captions). We then constructed voxelwise encoding models^24^ (Fig. 1a) to predict brain responses (fMRI) arising from *viewing* images from the Natural Scenes Dataset (NSD)^25^. We found that brain prediction performance was consistently higher for CLIP as compared to these other models.

**Figure 1.**
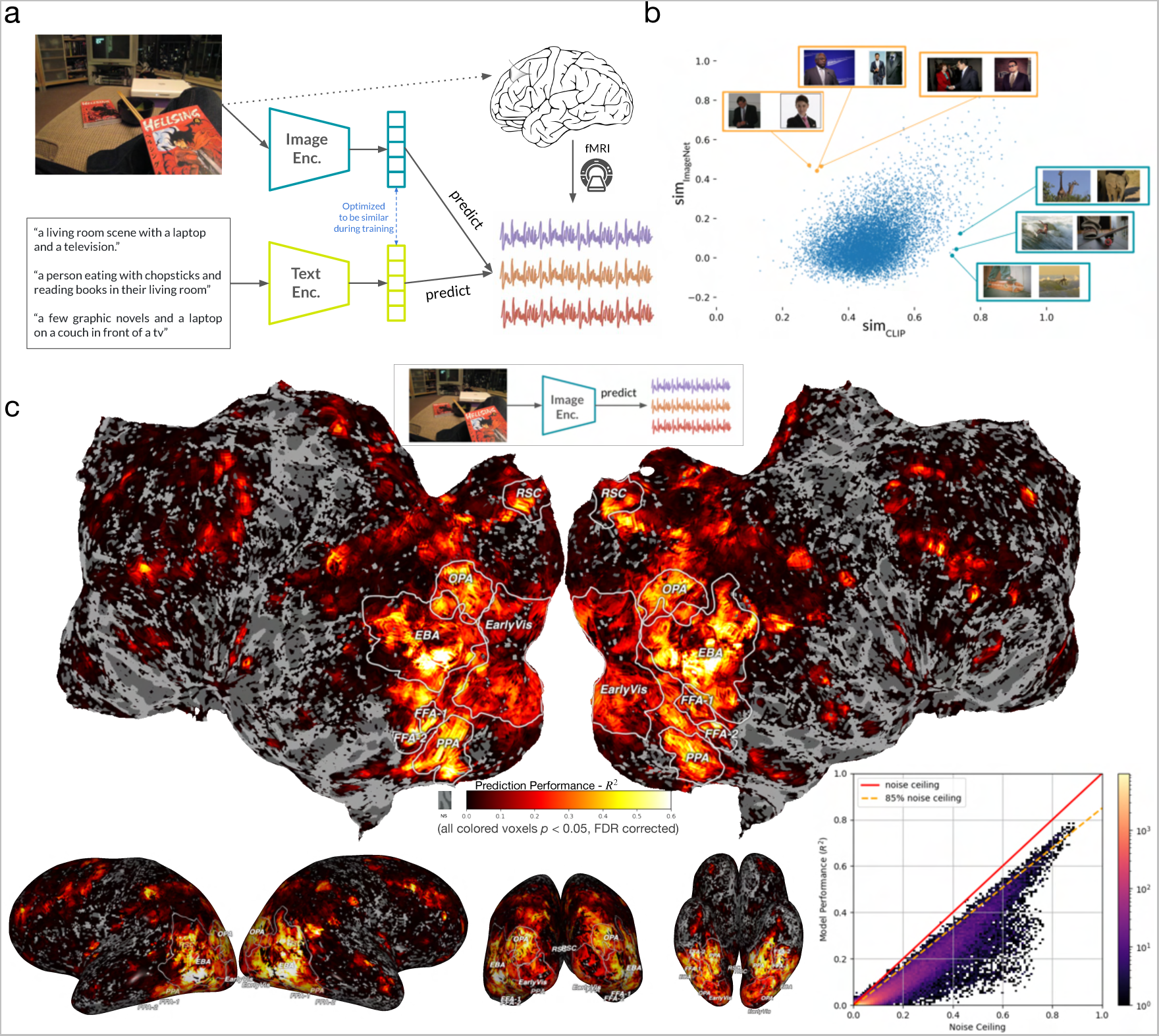
Model pipeline, motivation and prediction performance for the ResNet*_CLIP_* visual encoder. **(a)** Last-layer representations from the CLIP image and text encoders are extracted from images and captions, respectively. These representations are used in voxelwise encoding models to predict brain responses to each image. **(b)** The similarities of pairs of images when using embeddings from ResNet*_CLIP_*and ResNet*_ImageNet_* are compared. For each pair of 10000 randomly sampled images, a similarity score was computed (measured in correlation) between the representations of these two images extracted from ResNet*_CLIP_* and ResNet*_ImageNet_* (i.e. 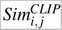 and 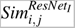). Pairs of images were ranked according to the differences between the similarities. Namely, 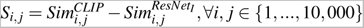, where 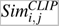 and 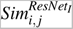 are correlations of representations between Image *i* and Image *j* in ResNet*_CLIP_* and ResNet*_ImageNet_*, respectively. The position of each dot in the scatter plot is determined by similarity scores for the same pair of images in ResNet*_CLIP_* and ResNet*_ImageNet_*model spaces. Pairs of images in the bottom right corner are those most similar in ResNet*_CLIP_* and most dissimilar in ResNet*_ImageNet_* (ranked by *S_i j_*); for example, images of people surfing and skateboarding and images of giraffes and an elephant. In contrast, pairs of images in the top left corner are those most similar in ResNet*_ImageNet_* and least similar in ResNet*_CLIP_*; for example, visually similar pictures of people wearing dark suits with a white shirt and a contrasting tie. **(c)** Voxelwise prediction performance (measured in *R*^2^) on a held-out test set is shown for Subject S5 in a flattened view of the brain with overlays for functionally-defined, category-selective ROIs (top), as well as in lateral, posterior and bottom views (bottom row, left-to-right). (Bottom-right) A 2D histogram of model performance in *R*^2^ against noise ceiling and 85% noise ceiling across all voxels in the whole brain. Density of voxels are shown in a log scale. Most voxels are predicted close to its noise ceiling and some are above the 85% noise ceiling. For visualization purpose, we only plot in the flatmap the voxels that are predicted significantly higher than chance (*p* < 0.05, FDR-corrected^29^).

A variety of factors characteristic of CLIP, and different from most prior models used for brain prediction, may be contributing to this superior prediction performance. However, as a proprietary model, we are unable to individually vary these factors. To explore these factors in a controlled manner, we selected several recent open source models that allow for more direct comparisons between four factors: architecture, feedback, dataset size, and data diversity. Our extended analyses included a self-supervised model, simCLR^26^, a self-supervised model that included language feedback, SLIP^17^, and several open versions of CLIP^27^. These models were trained with datasets that included 15 million (YFCC^28^), 400 million (as in the original CLIP model), or 2 billion examples from LAION^27^. We constructed encoding models with these networks to explain responses from NSD, which allowed us to more precisely evaluate and quantify the contributions of architecture, pre-training, dataset size, and data diversity.

To preview our findings, models using CLIP lead to encoding models that are much better at predicting high-level visual representations in the human brain as compared to single modality models that are pre-trained with smaller and less diverse datasets. Controlled comparisons suggest that these improvements are *not* due to architectural differences and, beyond a certain training dataset size, are related to the diversity of the data, as well as the joint image/caption training that this data affords. Critically, when dataset factors are controlled, we still see improvement with language feedback. We are also able to predict visual brain responses using image captions alone – indicating that models using CLIP learn a robust latent space bridging natural language and vision. In this vein, we observed the greatest improvements in prediction and were able to account for more of the unique variance in visual regions that process scenes of humans interacting with one another and their environment.

## Results

### Multimodal embeddings best predict high-level visual cortex

Our central question is whether incorporating joint natural language and image pre-training with large, diverse datasets leads to better models for understanding human high-level visual cortex. As a first step, we extracted representations from the last layer of the ResNet*_CLIP_* image encoder and ResNet*_ImageNet_* – networks that have the same architecture but are trained with different objectives. In a later section we address the fact that they were also trained with datasets that differ in size and diversity.

We expect that images are represented differently by ResNet*_CLIP_*and ResNet*_ImageNet_*, such that ResNet*_CLIP_* embeddings contain more semantic information and ResNet*_ImageNet_* embeddings contain more visual information. In Figure 1b, we show the pairwise similarity between embeddings of 10,000 randomly sampled stimulus images in both ResNet*_CLIP_* and ResNet*_ImageNet_*. Image representations in the two model spaces are correlated. However, when zooming in to the “corner” images in the similarity analysis (Fig. 1b), images represented more similarly in ResNet*_CLIP_*, but not in ResNet*_ImageNet_*, are semantically related, while those represented more similarly in ResNet*_ImageNet_* are visually related. Within ResNet*_CLIP_*, images of people surfing and skateboarding, as well as images of giraffes and an elephant, are more similar. In contrast, within ResNet*_ImageNet_*, images with different contexts are more similarly represented according to their visual similarity. For example, people wearing dark suits with a white shirt and a contrasting tie. These corner images indicate that ResNet*_CLIP_* captures contextual similarities that are not present in ResNet*_ImageNet_*.

We used the image representations from ResNet*_CLIP_*image encoder and ResNet*_ImageNet_* to predict fMRI voxelwise responses across the brain. In Figure 1c we show the *R*^2^ performance in the held out data across the whole brain. The overall level and the pattern of prediction performance were both highly consistent across S1-S8 (Fig. 1 for S5, Supplementary Figs. S2/S3 for S1–S8). The encoding model built with the last layer of ResNet*_CLIP_*’s visual encoder explains variance close to the voxel noise ceiling (see Supplementary Fig. S1 for performance measured in *r*). As a reference, earlier voxelwise encoding models for brain prediction report well below 0.7 in maximum correlation^30,31^, while in Allen et al.^25^, a brain optimized model of *early* visual cortex explains up to 0.8 in *R*^2^; similar to what we observe here in *high-level* visual cortex. However, directly comparing performance across models is challenging in that different studies use distinct experimental designs and rely on different data processing pipelines. Studies that report performance as averages within ROIs or as representation similarity (RSA) scores are also difficult to compare to our results.

Beyond overall performance metrics, peaks in the brain prediction maps were aligned with common category-selective ROIs. Peaks within regions implicated as scene-selective^32^, body-selective^33^, and face-selective^34,35^ were sufficiently well defined so as to allow ROI localization based solely on the prediction performance of ResNet*_CLIP_*. We speculate that these alignments reflect the importance of semantic associations in scene understanding and person recognition.

In order to rule out performance improvements based on specific network architectures, we extracted features from two backbones pre-trained with CLIP: visual transformer (ViT-32) and ResNet50. Differences in prediction performance were small (Supplementary Fig. S6), indicating that the observed improvement was not due to any particular neural-net architecture.

To explore whether the captions associated with images could predict the brain activity for viewing the corresponding image, representations extracted from the last layer of the CLIP text encoder were also used to predict voxelwise responses across the brain. In the absence of image information, we were able to predict responses in high-level visual cortex comparable to predictions from ResNet*_CLIP_*’s image encoder (Fig. 2). From this result, we infer that pre-training with CLIP enables learning a meaningful latent space that bridges between vision and natural language as is represented in the brain. At the same time, the text encoder explained less unique variance than the CLIP visual encoder, especially in the early visual cortex, suggesting that this is not a general effect (Supplementary Fig. S7).

**Figure 2.**
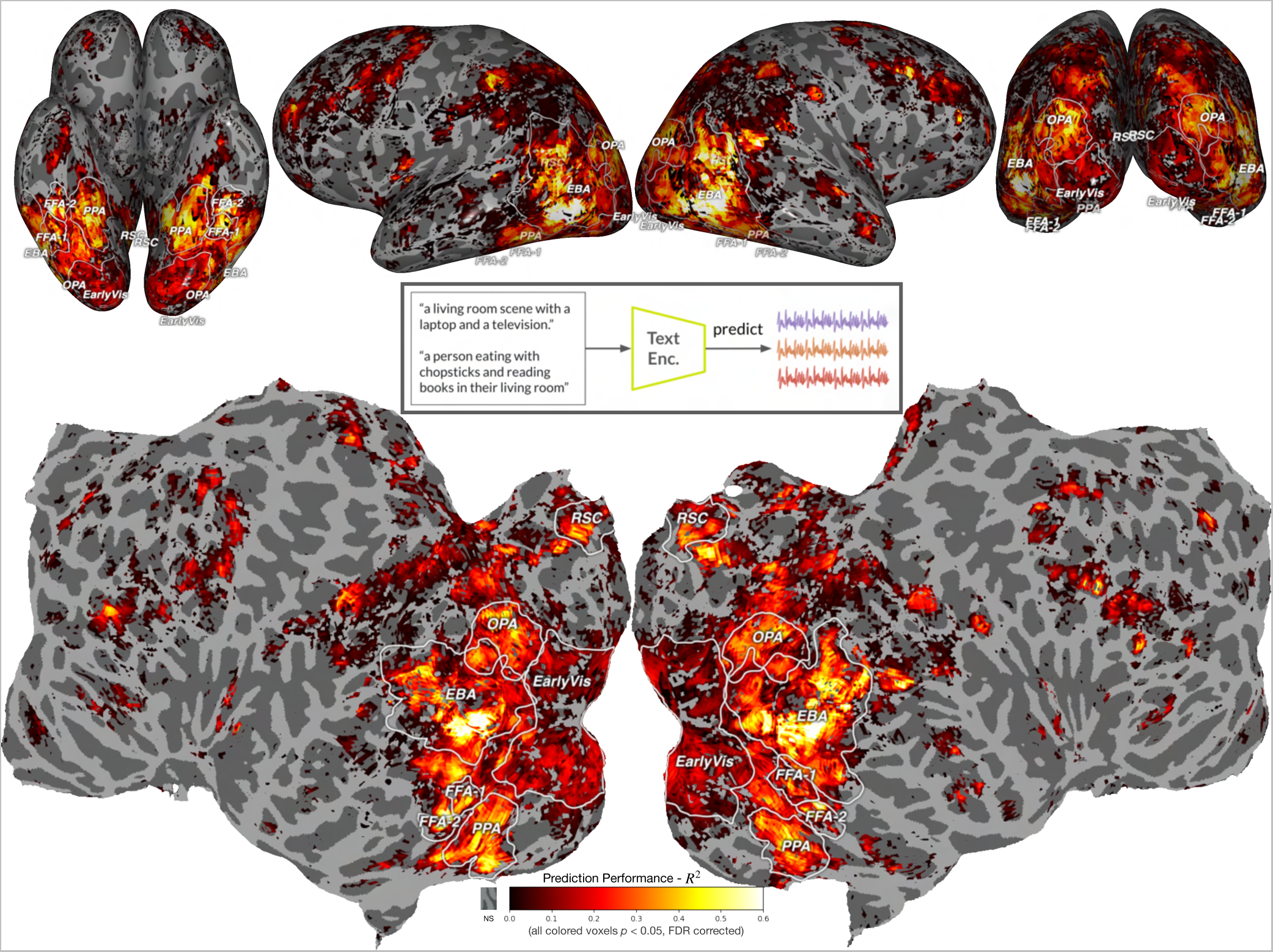
Prediction performance for the CLIP text encoder. Prediction performance for voxelwise responses – *R*^2^ – in held out data for the CLIP text encoding model for S5 with overlays for functionally-defined, category-selective ROIs. Captions of the images viewed in the scanner were provided to the text encoder and the representation was then used to make voxelwise brain predictions. Although only having access to the captions of the images that the subjects viewed, the CLIP text encoder was still able to predict fMRI data in many functionally-defined ROIs (e.g., EBA, PPA, RSC, FFA). The similarity in brain prediction for the image and text encoders reinforces the hypothesis that the information encoded in high-level visual areas is anchored in semantics.

### Visual embeddings with CLIP explain more unique variance than unimodal embeddings

To assess the impact of joint natural language and image pre-training with large, diverse datasets, we compared the unique variance accounted for by the last layer of ResNet*_CLIP_*’s image encoder to the last layer of ResNet*_ImageNet_* (with the same ResNet50 architecture).

A variance partitioning analysis^36,37^ (Fig. 3) revealed that ResNet*_CLIP_* accounts for the majority of the unique variance in high-level visual cortex, particularly in ROIs implicated in scene and person perception. With the exception of early visual areas (e.g., V1v, h4v), last layer of ResNet*_CLIP_* accounts for more of the unique variance for the majority of voxels in high-level ROIs (Fig. 3b). Beyond category-selective ROIs that respond to faces, places, and bodies, two areas, TPOJ and Angular Gyrus (AG), associated with theory of mind and language^38^, were likewise better explained by ResNet*_CLIP_*.

**Figure 3.**
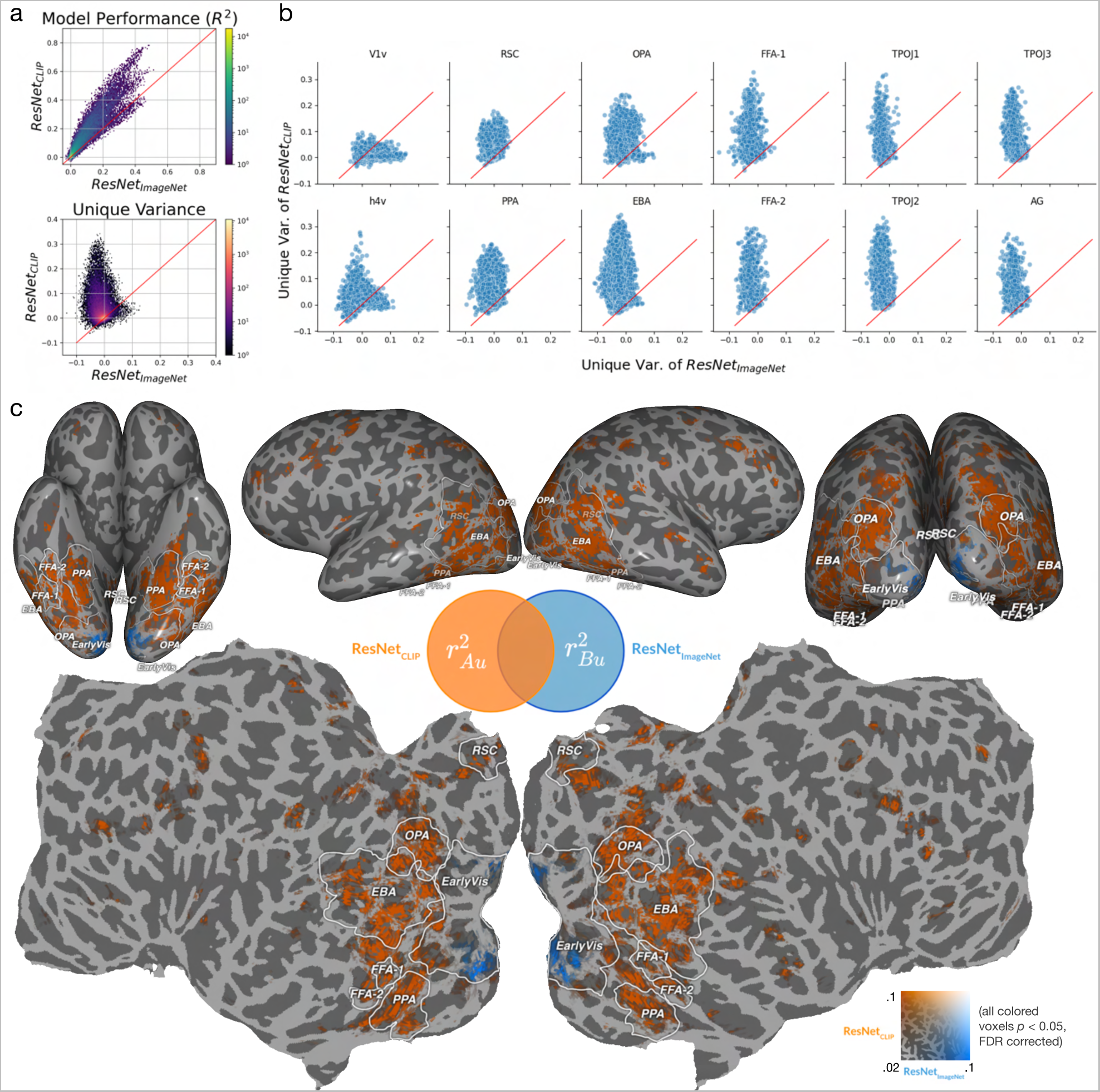
Performance for the CLIP visual encoder using a ResNet backbone as compared to ResNet*_ImageNet_*. **(a)** 2D distribution plots of voxels from the whole brain in S5 in model performance (in *R*^2^) and unique variance comparing between ResNet*_CLIP_* and ResNet*_ImageNet_*. The red lines indicates equal performance for the two models. ResNet*_CLIP_* predicts much better in terms of total variance and unique variance. **(b)** Unique variance accounted for by ResNet*_CLIP_* as compared to ResNet*_ImageNet_* for 12 different ROIs for all eight subjects. Individual voxels are plotted as blue points. The red lines indicate iso-variance, that is, (*y* = *x*). ResNet*_CLIP_* accounts for overwhelmingly more variance than ResNet*_ImageNet_* in high-level visual cortex. In contrast, ResNet*_ImageNet_* only accounts for more variance in ventral V1 and a reasonable proportion of the variance in ventral V4. **(c)** Unique variance accounted for by ResNet*_CLIP_* as compared to ResNet*_ImageNet_* for S5 – obtained by subtracting *R*^2^ for each model from that of the joint model (with concatenated feature spaces). Voxels where ResNet*_CLIP_* accounts for greater unique variance are orange and voxels where ResNet*_ImageNet_* accounts for greater unique variance are blue. Only voxels with significantly higher than chance unique variance are plotted for both models (*p* < 0.05, FDR-corrected).

The last layer of ResNet*_CLIP_* explained less variance in *early* visual cortex as compared to ResNet*_ImageNet_*; however, this does not imply that ResNet*_CLIP_* failed to capture information represented in these regions. The last layer of ResNet*_CLIP_* is the bottleneck layer that encodes image embeddings optimized to match in similarity with text embeddings. The entire visual pathway is best predicted by a progression of ResNet*_CLIP_* layers (including below the bottleneck layer; Supplementary Fig. S8). More generally, ResNet*_CLIP_* is the best predictive model for visual cortex.

### Regions that benefit most from ResNet***_CLIP_*** embeddings encode scenes of humans interacting with their environment

To explore the semantic dimensions learned in the encoding model built with CLIP, we performed principal component analysis (PCA) on the learned encoding model weight matrix concatenated across the 20,000 top predicted voxels from each of the eight subjects in NSD. We projected the concatenated voxels onto the principal component (PC) dimensions to understand the tuning of the entire voxel space, following previous works^31,39^. Visualizing each PC of the learned model and its corresponding voxel projection revealed the dimensions that underlie semantic organization in the brain. To interpret different PCs, we visualized the images that lie closest to a given PC in the model space for the top 5 PCs (which account for most of the explained variance; Supplementary Figs. S10/S11).

Through PC visualization, we found that PC1 separates animate and inanimate images and its brain projections correspond to body and face regions (Fig. 4d). PC2 separates scene and food images. When we split the functional areas identified from PC1 with PC2; its brain projections corresponded to place regions and the food region (Supplementary Fig. S12)^40–42^. We obtained interpretable PC dimensions up to PC10 (despite the relatively low explained variance from PC6 onwards), allowing us to identify fined-grained semantic distinctions (Supplementary Fig. S11).

**Figure 4.**
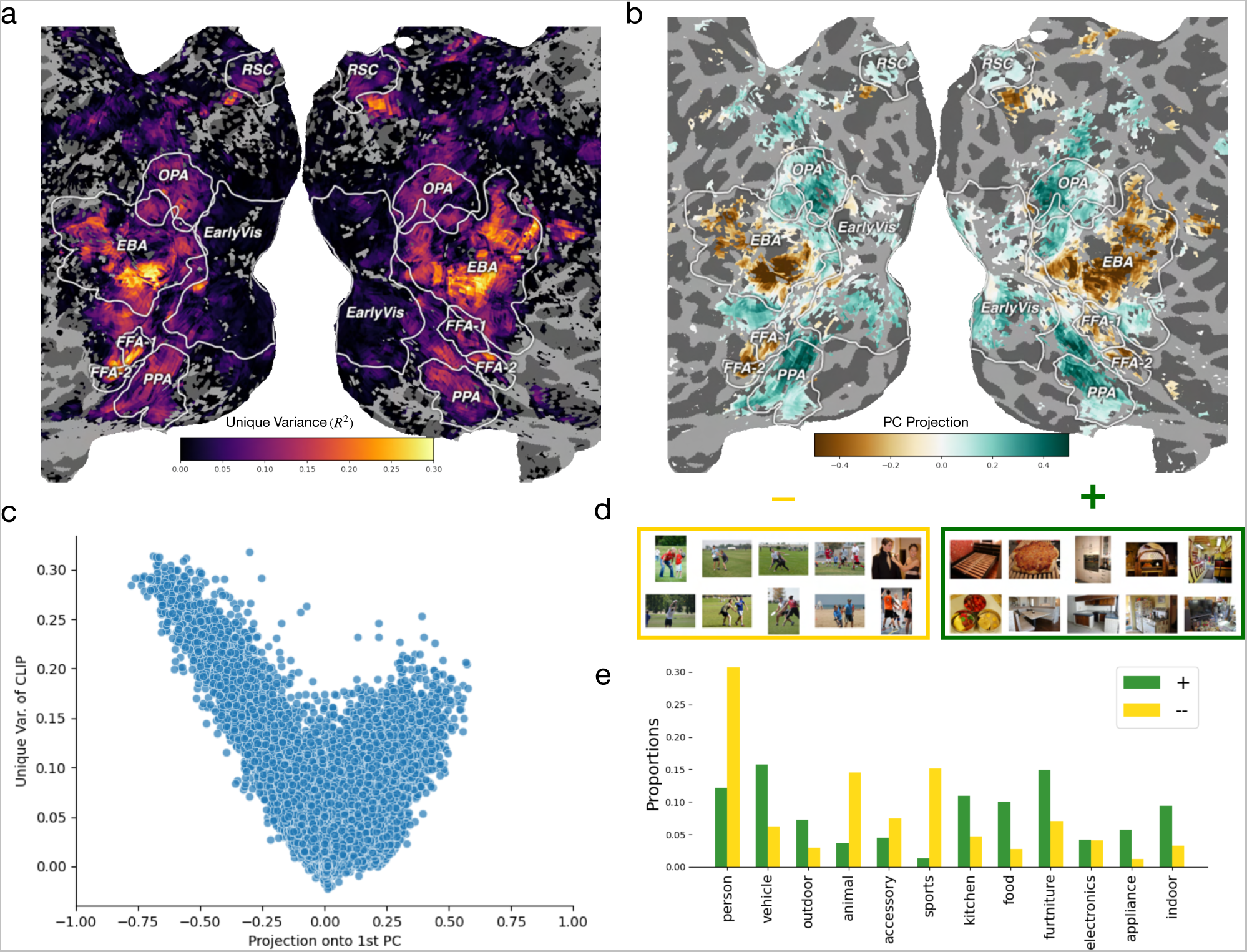
Better representations of scenes with people in a model trained with CLIP can account for gains in unique variance. **(a)** Unique variance explained by ResNet*_CLIP_* plotted on a flatmap from S5. **(b)** Projection of voxels onto PC1 of ResNet*_CLIP_*for S5. The voxels that are best explained by ResNet*_CLIP_* overlap largely with the voxels that lie on positive side when projected onto the 1st PC. **(c)** A voxelwise scatter plot illustrating that, for voxels lying on the negative side of 1st PC projection, the further down on the projection that the voxel lies, the better it is explained by ResNet*_CLIP_*. Note that the sign of the PC is arbitrary and can be flipped; we use “negative” here to refer to one of the sides of PC1. **(d)** Images are grouped into “+” and “−” depending on which side of the PC the image lies on when projected onto the PC1. The top 10 images that best align with either end of PC1 are shown in the yellow and green boxes respectively. For the positive projection we observe images of inanimate indoor scenes, whereas for the negative projection we observe images of people participating in animate outdoor sports. **(e)** The category distribution of the two image groups validates the observation that images on the negative side, relative to images on the positive side, consist more of people, animals, and sports.

Comparing the brain projection for PC1 and the unique variance map for ResNet*_CLIP_* we found that voxels that have large negative values on PC1 overlap with voxels where ResNet*_CLIP_*explains the most unique variance (Figs. 4a and 4b). These voxels clustered in ventral EBA, FFA-1, FFA-2, as well as ventral RSC. The more negatively a voxel lies along PC1, the more unique variance that can be explained by ResNet*_CLIP_* for this voxel(Figure 4c).

The projected images can be used to interpret which images had the largest benefit in brain prediction using ResNet*_CLIP_*. The images lying on the negative side of PC1 are people participating in sports (Fig. 4d). This separation is consistent with the best predicted voxels from ResNet*_CLIP_*being centered on the body area. Further validating this finding, images that lie on the negative end of the PC1 contain more people, animals, and sports items (based on the known category and super-category labels of COCO images; Fig. 4d). These observations suggest that the representation of people in ResNet*_CLIP_* is the domain for which the model provided the most leverage for predicting brain responses. More generally, ResNet*_CLIP_* is more effective at capturing scene semantics as compared to models trained with image/label pairs.

### Disentangling the effects of language feedback, model architecture, dataset size, and data diversity

Beyond natural language supervision, CLIP training uses larger training datasets and may have greater diversity as compared to earlier models – factors that may also contribute to CLIP’s effectiveness in brain prediction. To assess the contributions of these factors for prediction, we include three variance partitioning analyses using additional models as listed with their relevant characteristics in Figure 5a. These models allowed us to use the publicly available YFCC (“Yahoo Flickr Creative Commons”)^28^ and LAION^27^ datasets to control for dataset size and diversity. Both the YFCC and LAION datasets provide sufficient multimodal data to retrain CLIP with different and better controlled dataset parameters.

**Figure 5.**
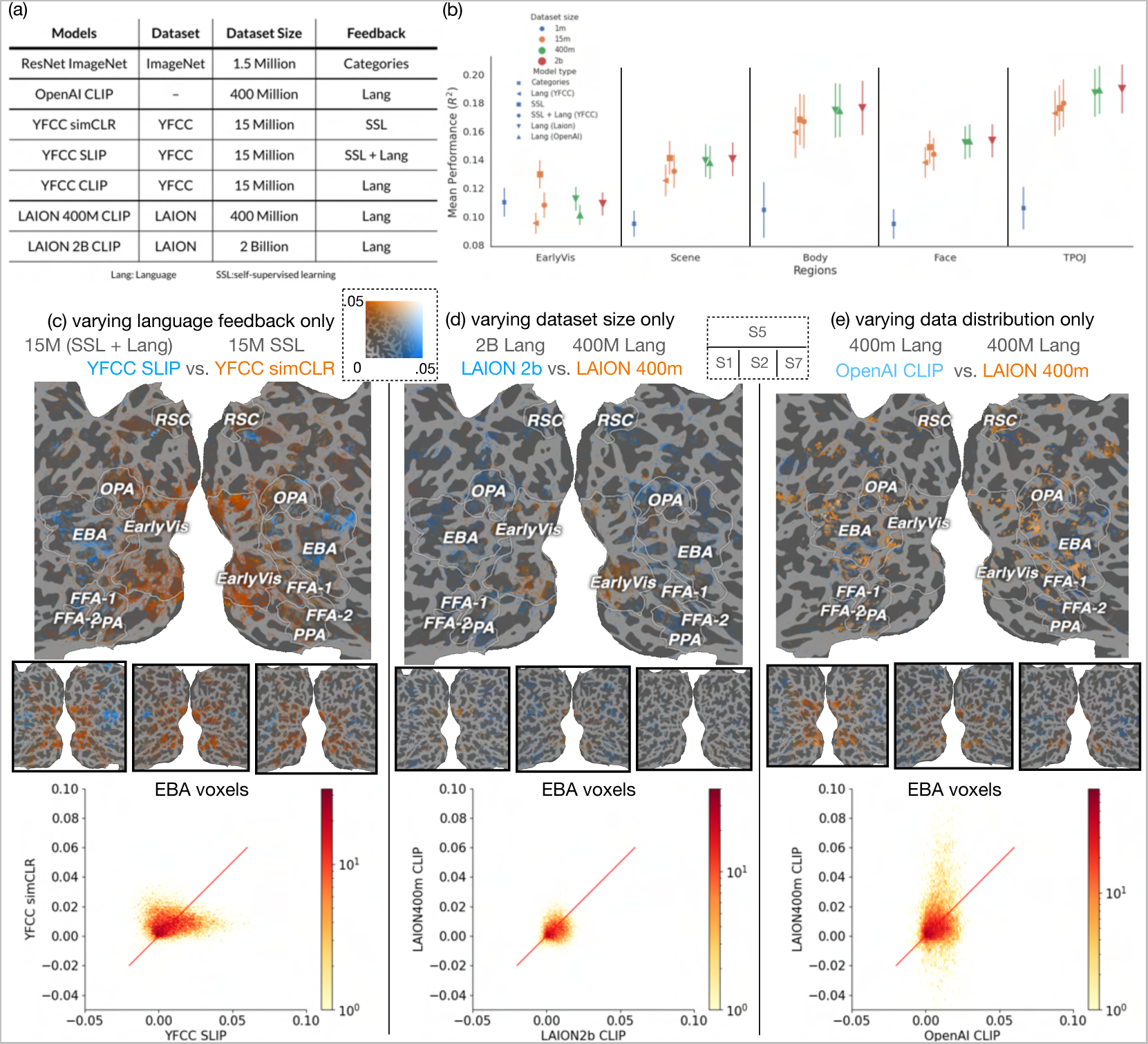
Variance partitioning analyses controlling for model architecture, data distribution, and dataset size indicate that dataset size and diversity have comparatively smaller effects on voxel prediction than language input. (a) The models we consider with their relevant characteristics; (b) Brain prediction performance averaged across all voxels in a given brain region for each model+dataset combination (“SSL” denotes self-supervised learning; “Lang” denotes natural language feedback for a given model). Each point is a region’s average performance across 8 subjects; error bars indicate standard error across 8 subjects. When looking at average brain prediction performance with an ROI, all three CLIP pre-trained models and the SSL model perform significantly better than ImageNet trained ResNet50, while differences between all three CLIP models and the SSL model are relatively small. (c-e) Cortical flatmaps showing the fine-grained, spatial distribution of unique variance for model comparisons varying a single factor while controlling for the others. Each unique variance comparison is thresholded by statistical significance (*p* < 0.05, FDR-corrected^29^). For each comparison, the first row shows brain maps from S5, while the second row shows unique variance brain maps from S1, S2, and S7, respectively. The third row of each comparison shows a 2D histogram of unique variance for individual voxels in EBA for all 8 subjects. The red line indicates identical unique variance (*y* = *x*). Notably, as shown in (c), when the same dataset is used for training across models, SLIP, a model that includes language feedback, accounts for more unique variance in high-level brain areas such as EBA and some parts of FFA, relative to simCLR, an otherwise identical model that does not include language feedback. Beyond language feedback, as shown in (e), a good data distribution appears to also account for unique variance in some high-level visual areas, while, as shown in (d), dataset size *per se* appears to account for very small improvements in unique variance past a certain size.

We visualized averaged model performance across all models in Figure 5b for several well-characterized ROIs within each general anatomical and semantic categories (EarlyVis: V1v, h4v; Scene: PPA, OPA, RSC; Body: EBA; Face: FFA-1, FFA-2; TPOJ: TPOJ-1, TPOJ-2). Each point in the figure is a region’s average performance across 8 subjects. All CLIP models and the SSL models explained brain responses in high-level visual cortex significantly better than ResNet*_ImageNet_*, while differences among SSL and CLIP are small. Note that these summary results describe *average* responses across all voxels in a given ROI, and therefore they do not reflect spatial patterns of unique variance within an ROI. Critically, ROI average analyses – particularly for ROIs containing large numbers of voxels – may mask meaningful spatial prediction patterns. For example, in work similar to ours, model comparisons that appear similar on average may actually carry fine-grained spatial information^43^.

To understand how model feedback, dataset size, and diversity affect predictions for individual voxels, we present cortical maps of the unique variance for three voxelwise analyses (Fig. 5c-e). Encoding models were used to predict individual voxel responses and the voxelwise unique variance explained by each model was computed.

We compared the effect of language feedback when controlling for the dataset parameters of distribution and size; the effect of dataset size when controlling for the data distribution, feedback type, and model architecture; and the effect of data distribution when controlling for feedback type, dataset size and model architecture. These three comparisons vary along single dimensions and, thus, are maximally informative in terms of isolating factors with respect to their impact on brain prediction performance. Other comparisons between these models would vary along more than one dimension, thereby confounding which factors were contributing to any observed effect.

We evaluated the effect of language feedback while controlling for dataset size and data distribution by comparing last layer of simCLR and SLIP (which combines simCLR and CLIP losses) trained on 15 million YFCC photo/caption pairs. We found more unique variance explained by SLIP in EBA, FFA and adjacent to the boundary of RSC, while simCLR showed more unique variance explained in early visual cortex and posterior EBA (Fig. 5c). The histogram of unique variance across all EBA voxels for all subjects revealed a bimodal distribution of voxels preferring one model or the other, with more voxels skewing towards YFCC SLIP. Flatmaps of significant unique variance explained by SLIP showed consistent patterns across subjects in MNI space (Supplementary Fig. S13). These visualizations suggest that interpreting brain prediction across models requires analysis at the voxel, rather than the ROI, level.

We evaluated the effect of dataset size while controlling for data distribution by comparing CLIP models trained on 400 million or 2 billion image/caption pairs from LAION^27^. The representations arising from the larger dataset explained more unique variance than those arising from the smaller dataset in EBA, FFA, and areas outside of RSC (Fig. 5d). However, the improvement in prediction performance due to dataset size was small. The histogram of EBA voxels revealed that dataset size, after reaching a critical level for model training, did not seem to be a major factor in improved brain prediction with CLIP.

We evaluated the effect of data distribution while controlling both feedback type and dataset size by comparing CLIP models trained on 400 million image/caption pairs from OpenAI^10^ and from LAION^27^. The representations using OpenAI’s dataset explained more unique variance than those using LAION’s dataset in regions including the EBA, FFA and areas outside of RSC (Fig. 5e). This aligns with the argument that data diversity in the training dataset contributes significantly to the robustness of the OpenAI CLIP model^16^. The histogram of EBA voxels revealed that differences arising from data distribution are larger than those arising from dataset size, indicating that data diversity is a likely factor in improved brain prediction with OpenAI CLIP.

## Discussion

Do higher performing models using natural language feedback together with larger, more diverse training sets also better predict brain response to complex, real-world scenes? We evaluated and quantified the contributions of large-scale, diverse multimodal pre-training as provided by CLIP for generating semantically-grounded representations of natural scenes. These models are extraordinarily good at predicting voxelwise responses to viewing scenes in the Natural Scenes Dataset^25^. A recent study confirms our results, finding that models with CLIP pre-training better predicted responses in NSD as compared to other models^43–45^.

While it is appealing to attribute the improved prediction performance of CLIP to language feedback during training, it is important to disentangle several other factors that may contribute to the high level of brain prediction performance. CLIP models, as compared to prior models used for brain prediction^2,30^, can have different model architectures and are pre-trained with many more examples. Consequently, model architecture, multimodal pre-training, dataset size, and data diversity (or some combination therein) are all characteristics of CLIP that may underlie improved brain prediction.

First, we found that model architecture, as realized in different visual backbones, had little impact on prediction performance (see also^43–45^). Second, to better isolate the potential contributions of other factors, we examined prediction performance for pairs of open CLIP models that differ from one another along only single dimensions. We found that: models trained with natural language feedback show a consistent advantage in prediction performance in high-level brain regions, especially the EBA and TPOJ; the diversity and quantity of the training data may set a ceiling for improvements in prediction arising from adding language feedback; the size of the training dataset for the CLIP model appears to be both less consequential for improved prediction performance and shows diminishing returns as compared to other data characteristics (i.e., data diversity). We conclude that models trained with natural language feedback, together with sufficient data diversity and dataset size, are attractive candidate models for understanding representation in high-level human visual cortex.

More broadly, rather than simply quantifying overall brain prediction performance across a range of models, we selected models with characteristics that reflect human-like training and experience in the form of natural language feedback and larger, more diverse datasets. In addition to ROI-level analyses, we also provided finer-grained analyses that shed light on how language training in tandem with a diverse dataset facilitates learning brain-like representations. Visualizations of ResNet*_CLIP_*and unimodal network representations revealed that ResNet*_CLIP_*better captured semantic information – consistent with natural language feedback being an important factor in improved prediction performance, as well as this improvement being associated with better prediction of high-level visual cortex. PCA analyses built on these results, revealed that, within ResNet*_CLIP_*, the fine-grained representation of scenes depicting human interactions lead to the largest gains in brain prediction – particularly in EBA. We suggest that ResNet*_CLIP_* captures information about humans interacting with the world around them, and, as such, is predictive of similar representations in high-level brain regions^46^.

Our results support the theory that, beyond object identity, human high-level visual representations reflect semantics and the relational structure of the visual world; for example, non-perceptual associations such as function or meaning^7,46,47^. An embedding model based on text captions for viewed images also concluded that high-level visual cortex represents semantic information related to those images^48^. Similarly, a study combining a deep neural network for vision with an attractor network of semantics was able to account for patterns of activation in high-level visual cortex^49^. Indeed, there is evidence that, along with playing a role in the acquisition of semantics^50,51^, language influences the acquisition of *visual* categories during development, where visual learning occurs concurrently with language and conceptual learning^51–53^. Our present study reinforces these findings, suggesting that language and semantics strongly influence the high-level organization of visual information encoded in the human brain.

What it is about models incorporating natural language training together with large, diverse datasets that enables them to excel not just at visual tasks such as zero-shot learning, but also at brain prediction? In our view, the same natural language feedback, in tandem with data diversity and dataset size, underlies higher performance in both domains – an example of the principle whereby higher performing models also perform better at brain prediction^3^. However, while dataset size may be a contributing factor to higher performance, beyond a certain size, its impact on brain prediction may be diminished except to the extent that size contributes to diversity. Our intuition is that re-training ResNet with a much bigger dataset, but continuing to include only category labels, would produce a model that would be unlikely to learn fine-grained representations of complex scenes. Such nuanced information regarding human interactions in real-world scenes is not carried by category labels: while some labels do contain semantic information beyond the category (e.g., “party”), language feedback provides context and a broader understanding, for example, the semantic relationships between actors and objects in real-world scenes. Supporting this point, given equivalent training data, models with natural language feedback outperformed self-supervised and unimodal models in high-level visual areas. Thus, the natural language feedback present in models pre-trained with CLIP appears crucial to their excellent performance in tasks related to both machine and biological intelligence.

In sum, the impressive ability of vision models incorporating natural language supervision along with large, diverse datasets for predicting brain responses opens new possibilities for developing a deeper understanding of the functional architecture of the human brain. Exploring the implications of this finding will require new ways of thinking about both artificial and natural systems. Future large-scale efforts should incorporate stimuli, tasks, representations, models, and datasets that reflect the natural complexity of how we interact with the world around us.

## Materials and Methods

### Datasets

#### fMRI data

Brain recordings were obtained from the the Natural Scenes Dataset (NSD)^25^, an open dataset of 7T whole brain high-resolution fMRI responses from eight subjects (S1-S8) who each viewed ~10,000 unique images of natural scenes, each image repeated 3 times. These scene images were a subset of the images in the annotated Microsoft Common Objects in Context (COCO) dataset^54^. Of the 70,566 total images presented across subjects, ~1,000 images were viewed by all subjects. fMRI data were collected during 30-40 scan sessions. Stimulus images were square cropped, presented for 3 s at a size of 8.4*^◦^* × 8.4*^◦^* with 1 s gaps in between image presentations. Subjects were instructed to fixate on a central point and to press a button after each image if they had seen that image previously.

The functional MRI data were acquired at 7T using whole-brain gradient-echo EPI at 1.8-mm resolution and 1.6-s repetition time. Preprocessing steps included a temporal interpolation (correcting for slice time differences) and a spatial interpolation (correcting for head motion). Single-trial beta weights were estimated with a general linear model. In this paper we used the betas_fithrf_GLMdenoise_RR preparation of the betas. FreeSurfer^55,56^ was used to generate cortical surface reconstructions to which the beta weights were mapped. The beta weights were z-scored across run and were averaged across repetitions of the image (up to 3 repetitions of each image), resulting in one averaged fMRI response to each image per voxel, in each subject. NSD also includes several visual ROIs that were identified using separate functional localization experiments. We drew the boundaries of those ROIs for each subject on their native surface for better visualization and interpretation of the results (e.g., Fig. 1). All brain visualizations were produced using Pycortex software^57^.

#### Natural scene images

All stimulus images used in NSD and in our experiments were drawn from the COCO dataset^54^. COCO is unique among large-scale image datasets in that COCO images contain contextual relationships and non-iconic (or non-canonical) object views. In comparison to ImageNet^5^, COCO contains fewer labeled categories (91), but includes more examples for each category (*>* 5, 000 for 82 of the categories). Note, however, that many labeled categories in ImageNet are at the subordinate level – COCO likely contains at least as many *unlabeled* subordinate categories. The complete set of COCO images and additional details can be found on the COCO website: https://cocodataset.org.

### Model details and feature extraction

Models used in the analysis includes: 1) OpenAI trained CLIP (with ViT-32 transformer and ResNet50 backbones); 2) YFCC trained SLIP, CLIP, simCLR; 3) Open CLIP models trained on LAION 400m and LAION 2B; 4) ImageNet pretrained ResNet50. YFCC is a 100 million example dataset comprised of multimedia “objects” which includes 15 million photos with captions selected from Flickr^28^, while LAION is a large-scale dataset that contains 5.85 billion multilingual CLIP-filtered image-text pairs^27^. All NSD stimulus images were input into the these models.

For model comparison, we use the output of the “image encoder” in CLIP models and the second to the last layer in ImageNet trained models as feature spaces for the encoding models. The feature dimensions for each of the model feature spaces are as follows: ImageNet trained ResNet50: 2048; OpenAI CLIP with ViT-32 backbone: 512; OpenAI CLIP with ResNet50 backbone: 1024; YFCC simCLR: 768; YFCC SLIP: 512; YFCC CLIP: 512; LAION 400m CLIP: 512; LAION 2B CLIP: 512.

For image captions, we use the human generated captions for each of the NSD images provided by the COCO dataset and input them into both BERT and CLIP-based models’ text encoders for their layerwise activations. On average, COCO provides 5-6 captions for each image. Caption embeddings for a image are extracted individually and the average is used in the encoding models.

### Voxelwise encoding models

We built ridge regression model (implemented in PyTorch; see^58^) to predict one averaged fMRI response to each image per voxel, in each subject. We chose to use a ridge regression model instead of more complicated models in order to retain the interpretability of model weights, which may provide insights into the underlying dimensions of the brain responses. We randomly split the total number of images a subject sees into training and test set with a 4-to-1 ratio. For each subject, each voxel’s regularization parameter was chosen independently via 7-fold cross-validation across the training set. We swept through 100 regularization parameters spaced evenly on a log scale from 10*^−^*^8^ to 10^10^, i.e. np.logspace(−8, 10, 100). Cross-validation was handled by `sklearn.model_selection.KFold`, where data are split into consecutive folds without shuffling. Each fold is used once as validation while the rest of the set are used for training. Model performance was evaluated on the test data using both Pearson’s correlation and coefficient of determination (*R*^2^). To determine the significance of the predictions, we perform a bootstrap test where we resample the test set with replacement for 2000 times and compute the FDR corrected *p*-values threshold for various performance statistics^29^.

### Variance partitioning

To obtain unique variance by two model A and B, we first create joint model of A and B by concatenating features from these two models. We then fit voxelwise ridge regression model to the joint model and obtain 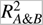. The variance explained by individual model A and B are denoted as 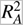 and 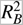, respectively. We then calculated the unique variance for model A and B, where 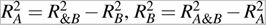.

### PCA analysis

To recover the semantic basis of the learned encoding model we performed principal component analysis (PCA) on the learned weight matrix concatenated across the 20,000 top predicted voxels from each of the eight subjects in NSD. These voxels were selected based on the noise corrected model performance of the ResNet*_CLIP_*. We then concatenated the weight matrices (used in encoding model with ResNet*_CLIP_*) corresponding to these voxels from all eight subjects along the voxel dimension. We then centered the matrix, applied PCA, and obtained the first 20 principal components (PCs). As in prior work^31,39^, we projected the concatenated voxels onto the PC dimensions to understand the tuning of the entire voxel space. Explained variance by these PCs are plotted in Supplementary Figure S10.

## Statistics and Reproducibility

Statistical analyses were performed using Python and data visualizations were accomplished using Pycortex**^?^**. Significant voxels from encoding models were identified by computing a *p*-value from each *R*^2^ and corrected for multiple comparisons using the the Benjamini-Hochberg False Discovery Rate procedure (FDR)^29^ and *α* = 0.05. Additional information on the analyses are provided in *Methods*.

## Data Availability Statement

The NSD data was made available by Allen et al.^25^. All other data are available from the corresponding author on reasonable request.

## Code Availability Statement

Our code is available as a public Github repository https://github.com/ariaaay/clip2brain.git.

## Acknowledgements

AYW and MJT were supported by AFRL/AFOSR award FA9550-18-1-0251. The NSD was supported by NSF IIS-1822683 and NSF IIS-1822929. We thank the following people for contributing technical assistance, ideas and commentary to this project: Jayanth Koushik, Nadine Chang, and Maggie Henderson.

## Author Contributions Statement

AYW, MJT, and LW conceived the experiments, KK and TN collected the neuroimaging data, AYW conducted the experiments, AYW analysed the results. All authors wrote and edited the manuscript.

## Author Declaration

The authors declare no competing interests.

---

**Supplementary Figure S1.**
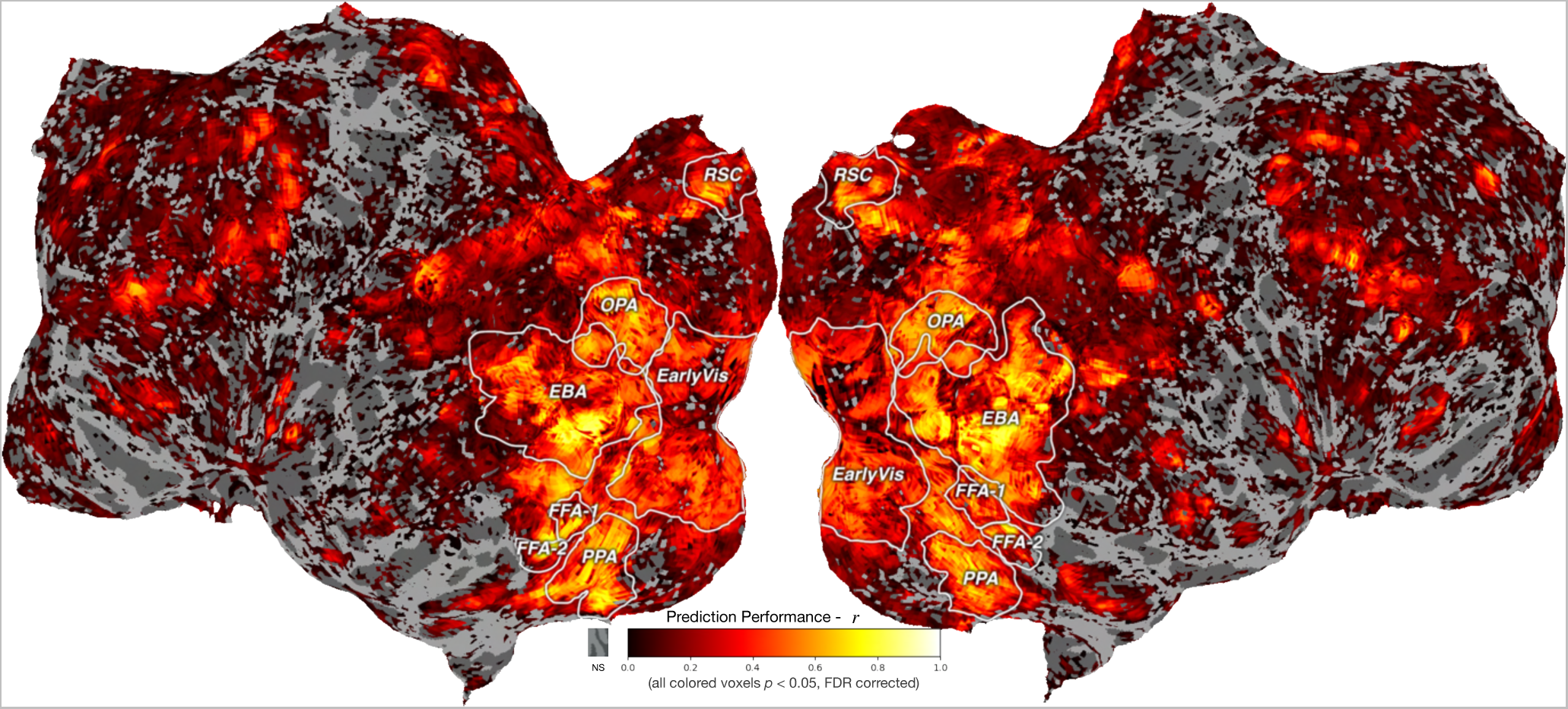
Prediction performance meansured in corrlation using the CLIP visual encoder. Voxelwise prediction performance (measured in *r*) on a held-out test set is shown for S5 in a flattened view of the brain.

**Supplementary Figure S2.**
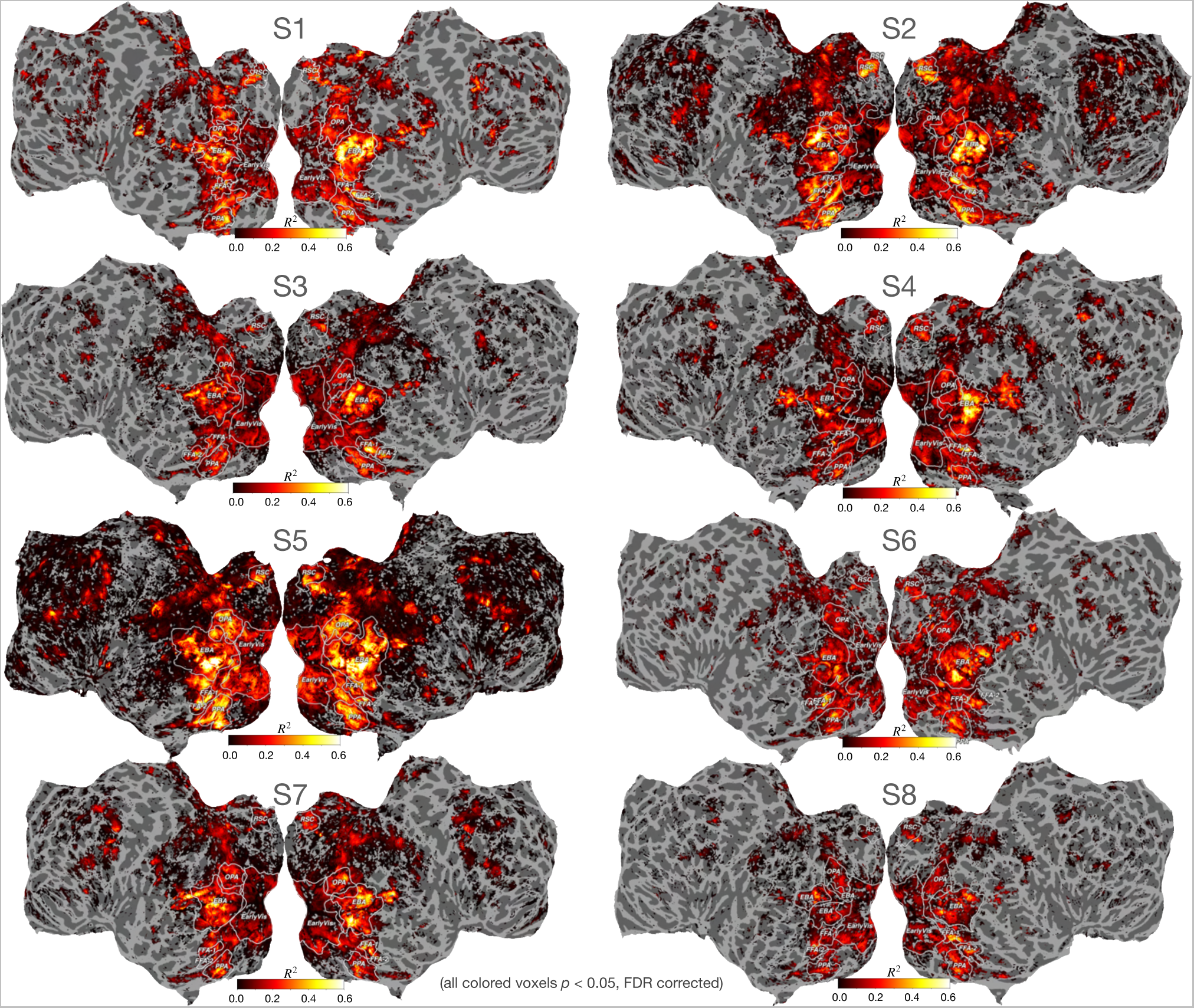
Prediction performance with CLIP visual encoder for all eight subjects. Voxelwise prediction performance (measured in *R*^2^) on a held-out test set is shown for S1-S8 in a flattened view of the brain.

**Supplementary Figure S3.**
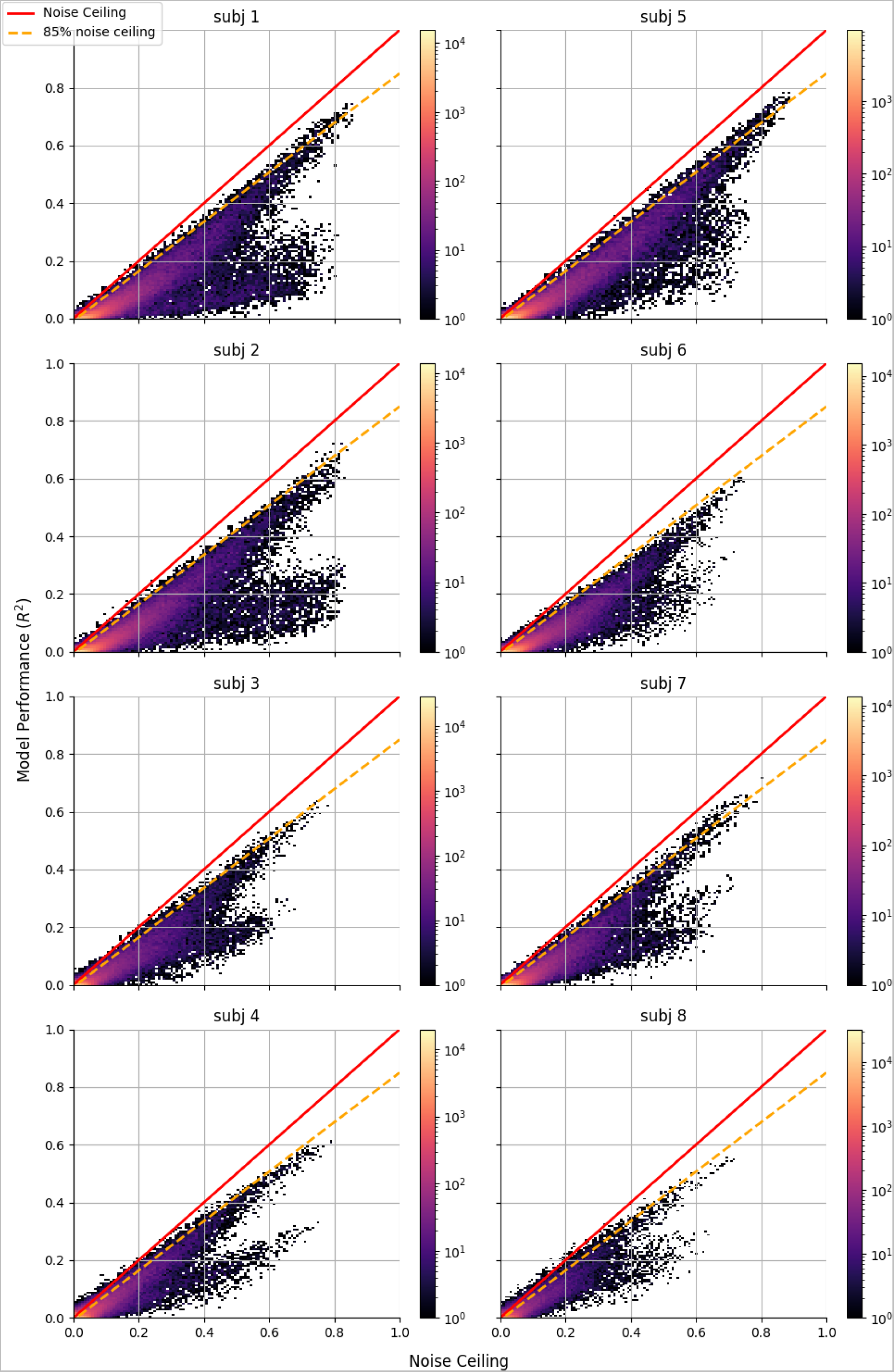
Scatterplots of noise ceiling against model performance in *R*^2^ for all subjects.

**Supplementary Figure S4.**
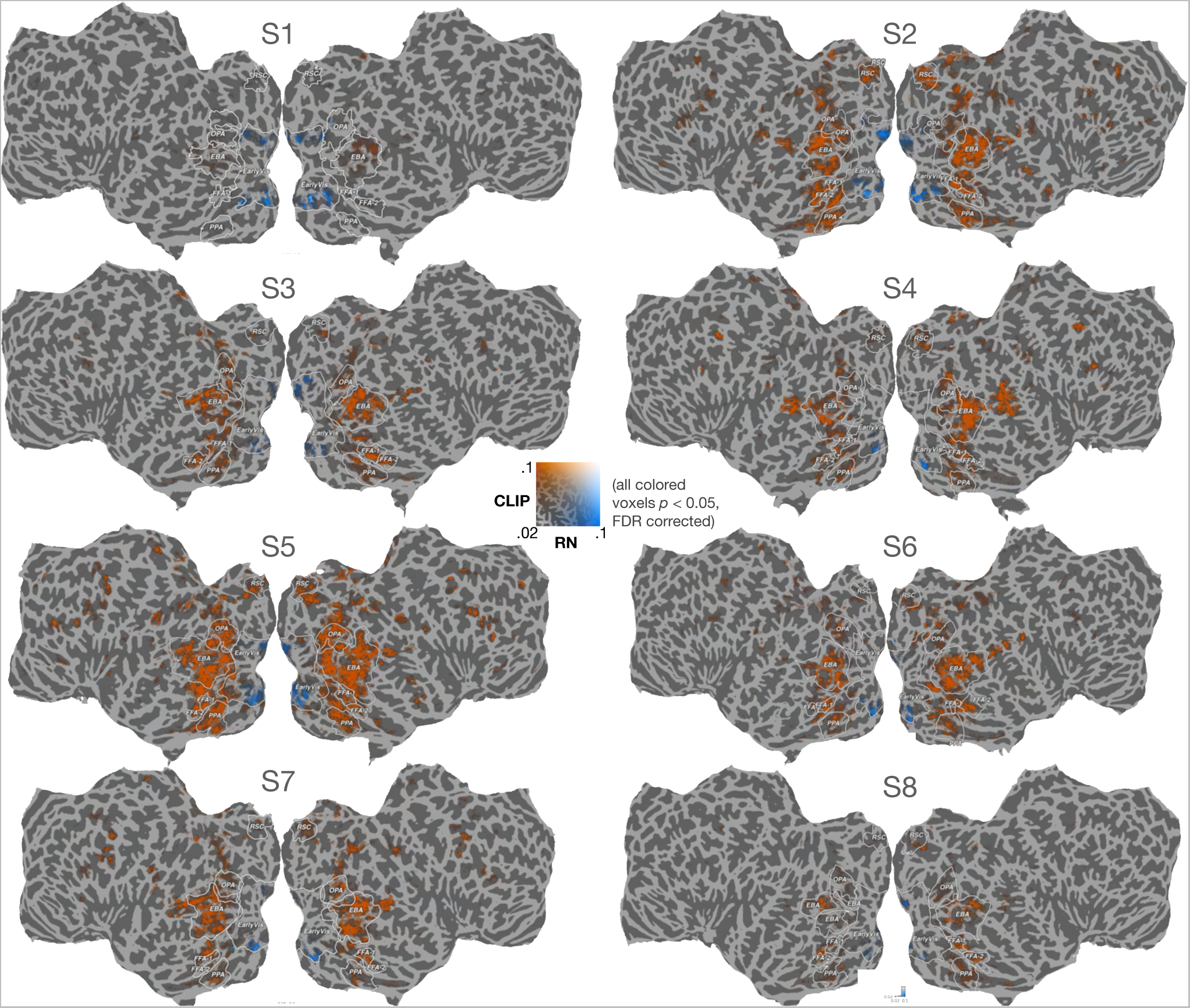
Unique variance accounted for by ResNet*_CLIP_* as compared to ResNet*_ImageNet_* (noted as RN in the figure) for all eight subjects. Voxels where ResNet*_CLIP_* accounts for greater unique variance are orange and voxels where ResNet*_ImageNet_* accounts for greater unique variance are blue.

**Supplementary Figure S5.**
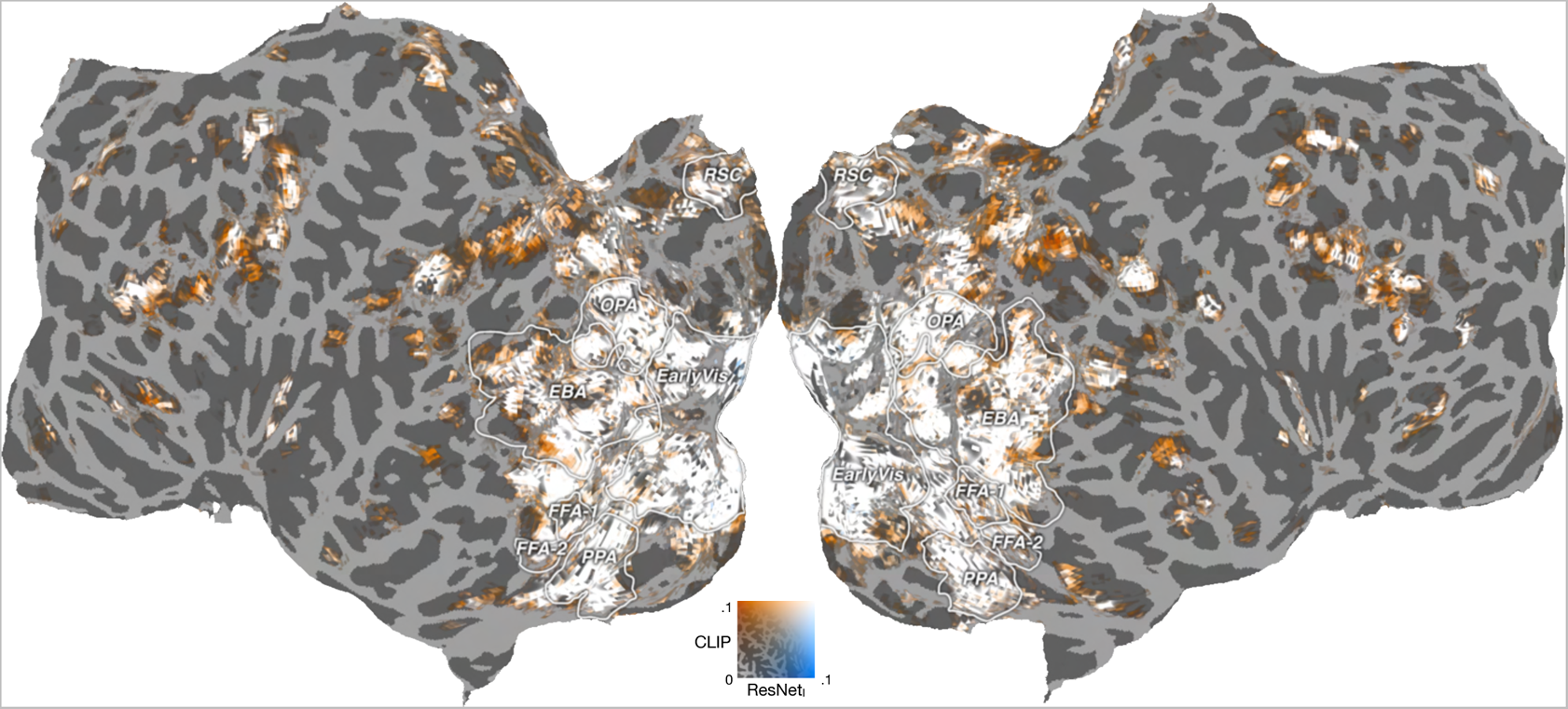
Total variance accounted for by ResNet*_CLIP_* as compared to ResNet*_ImageNet_* for S5. Voxels where ResNet*_CLIP_* accounts for greater variance are orange and voxels where ResNet*_ImageNet_* accounts for greater variance are blue. White voxels are where both models explain well.

**Supplementary Figure S6.**
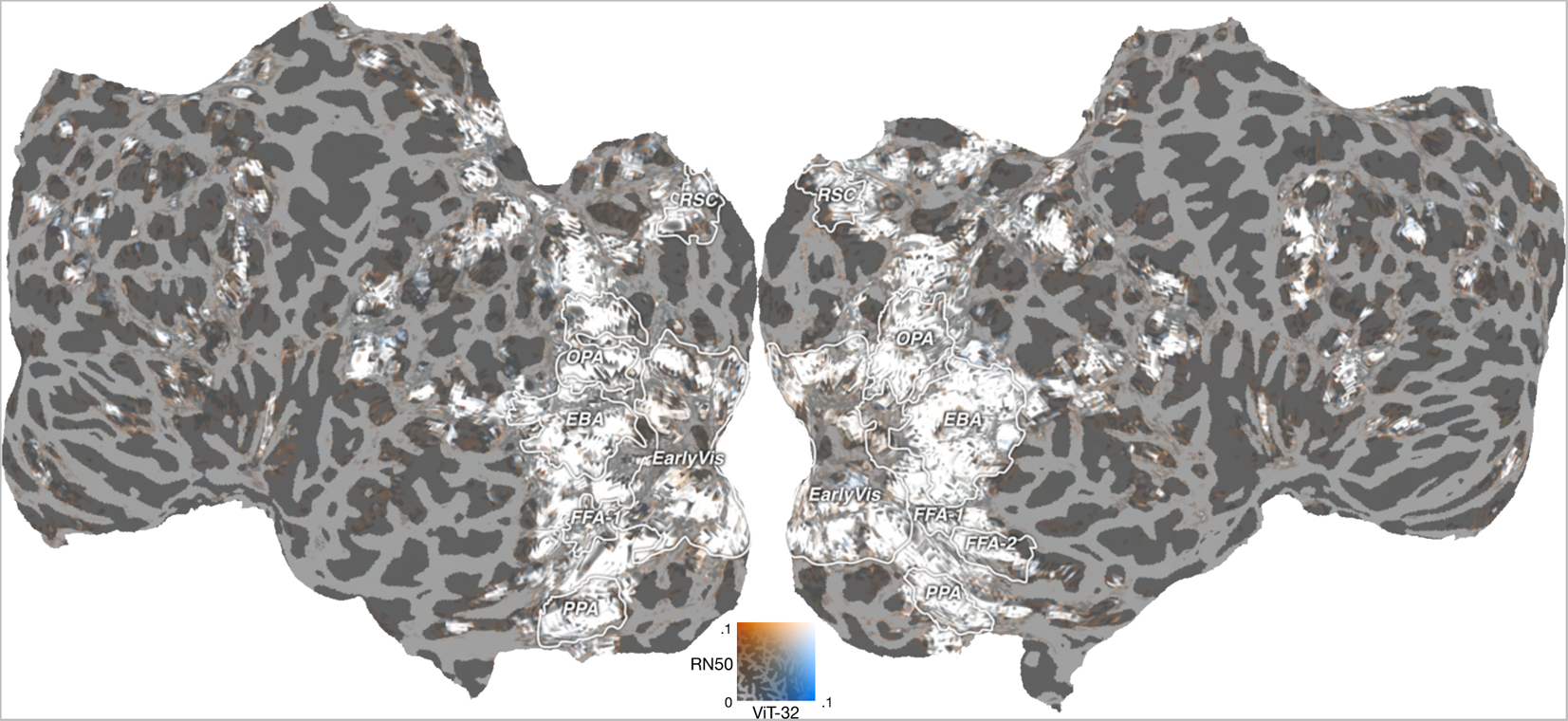
Performance 2D map between ResNet*_CLIP_* and ViT-32*_CLIP_*.

**Supplementary Figure S7.**
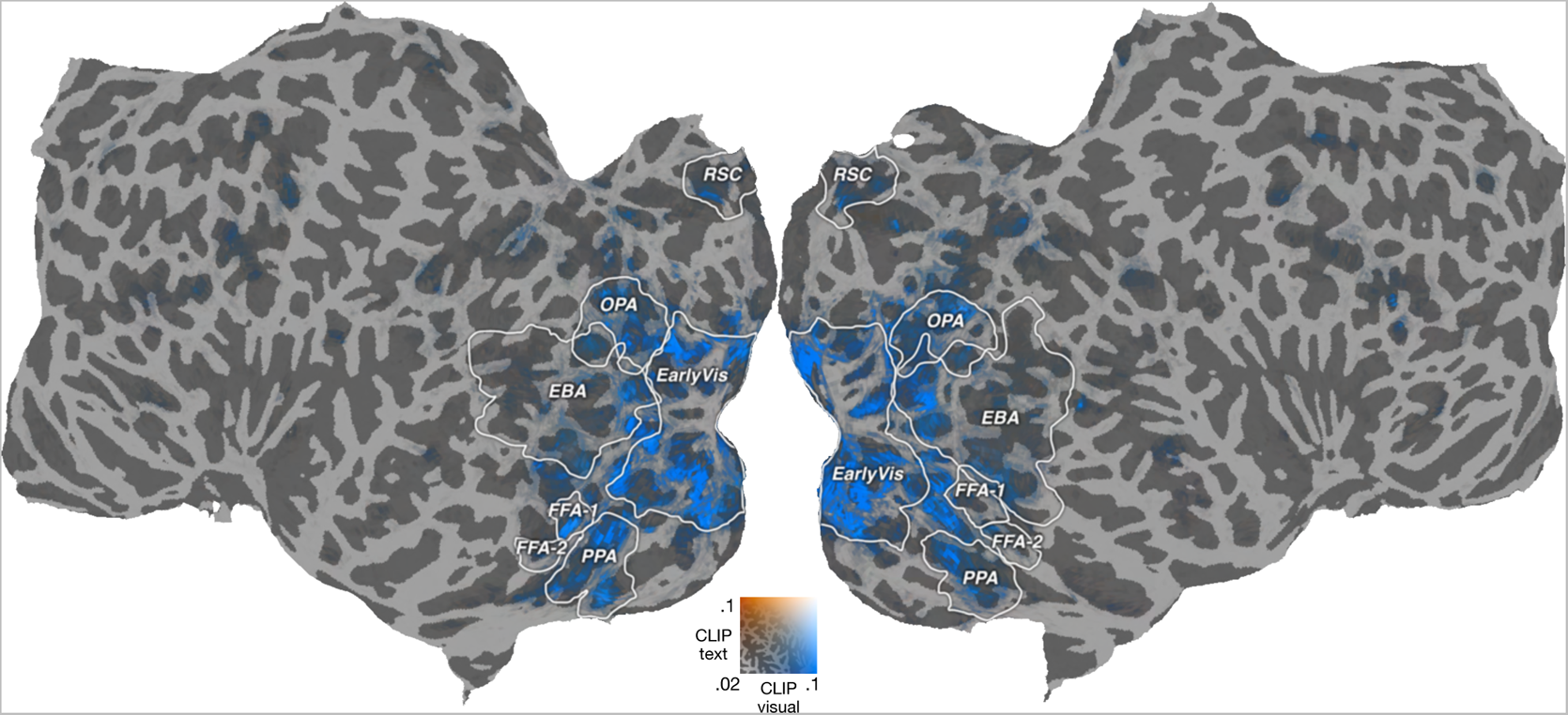
Unique variance by CLIP visual encoder and CLIP text encoder.

**Supplementary Figure S8.**
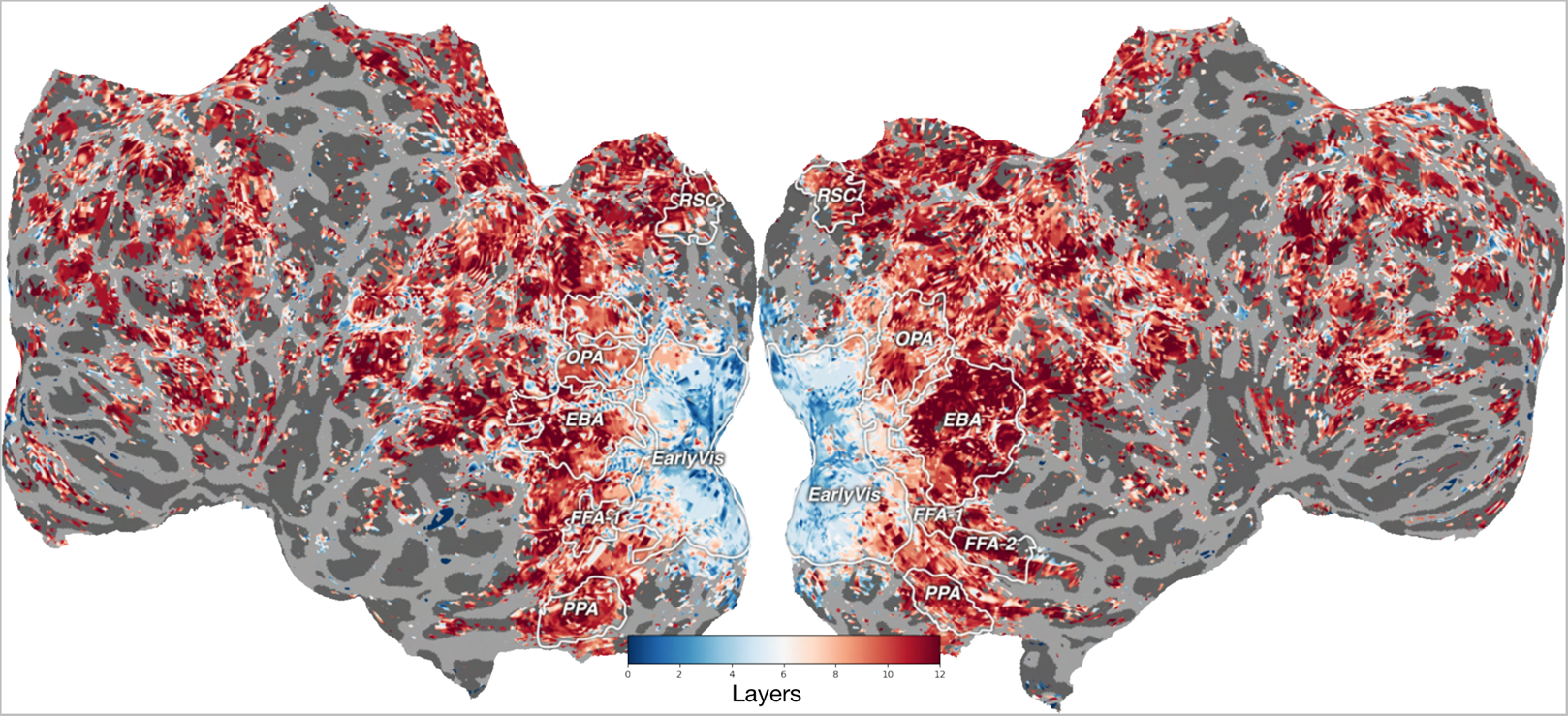
Layer preference by voxels across the brain.

**Supplementary Figure S9.**
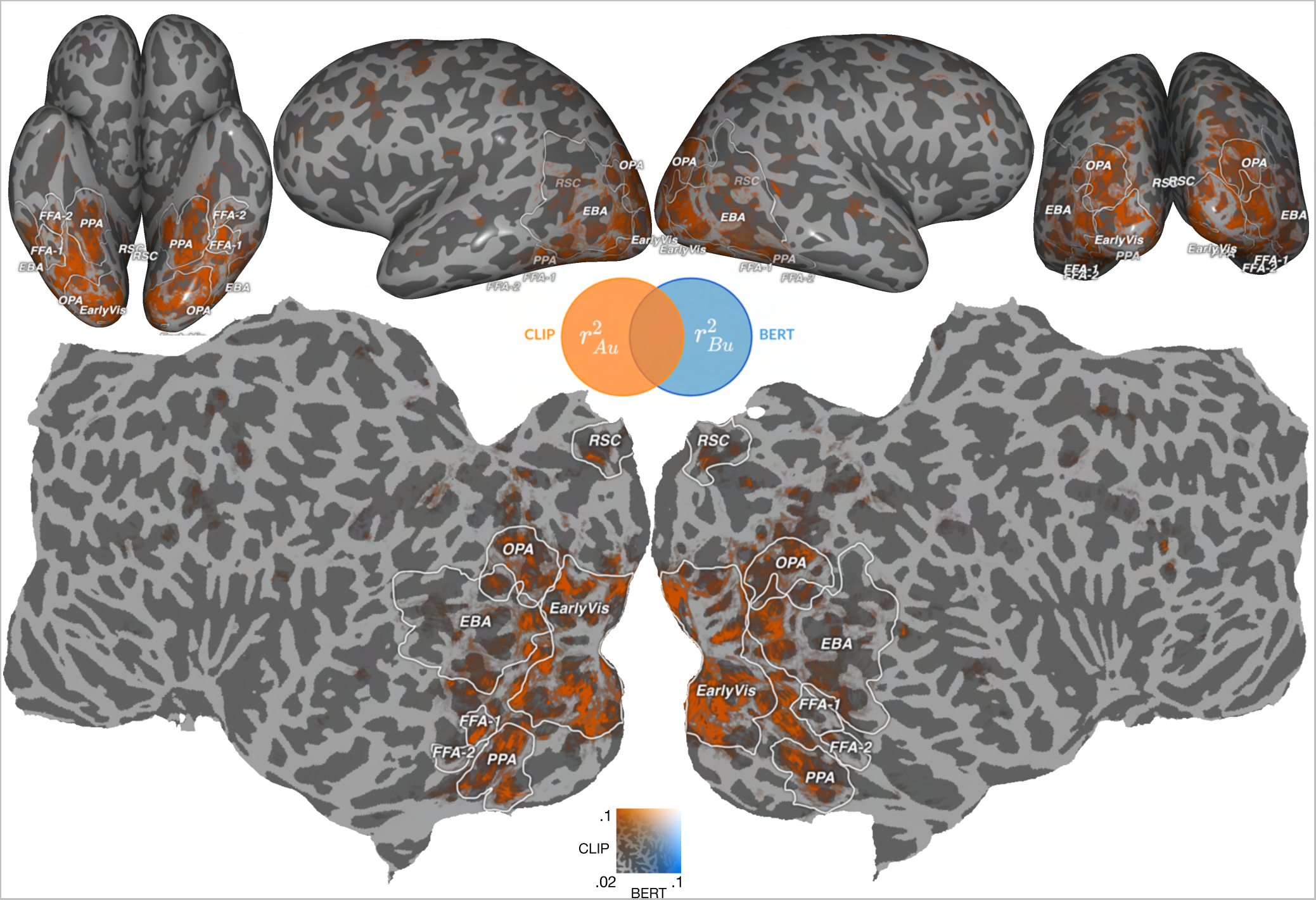
Performance comparison between CLIP text encoder with BERT. Unique variance accounted for by CLIP as compared to BERT for S5 – obtained by subtracting *R*^2^ for each model from that of the the concatenated model. Voxels where CLIP accounts for greater unique variance are orange and voxels where BERT accounts for greater unique variance are blue.

**Supplementary Figure S10.**
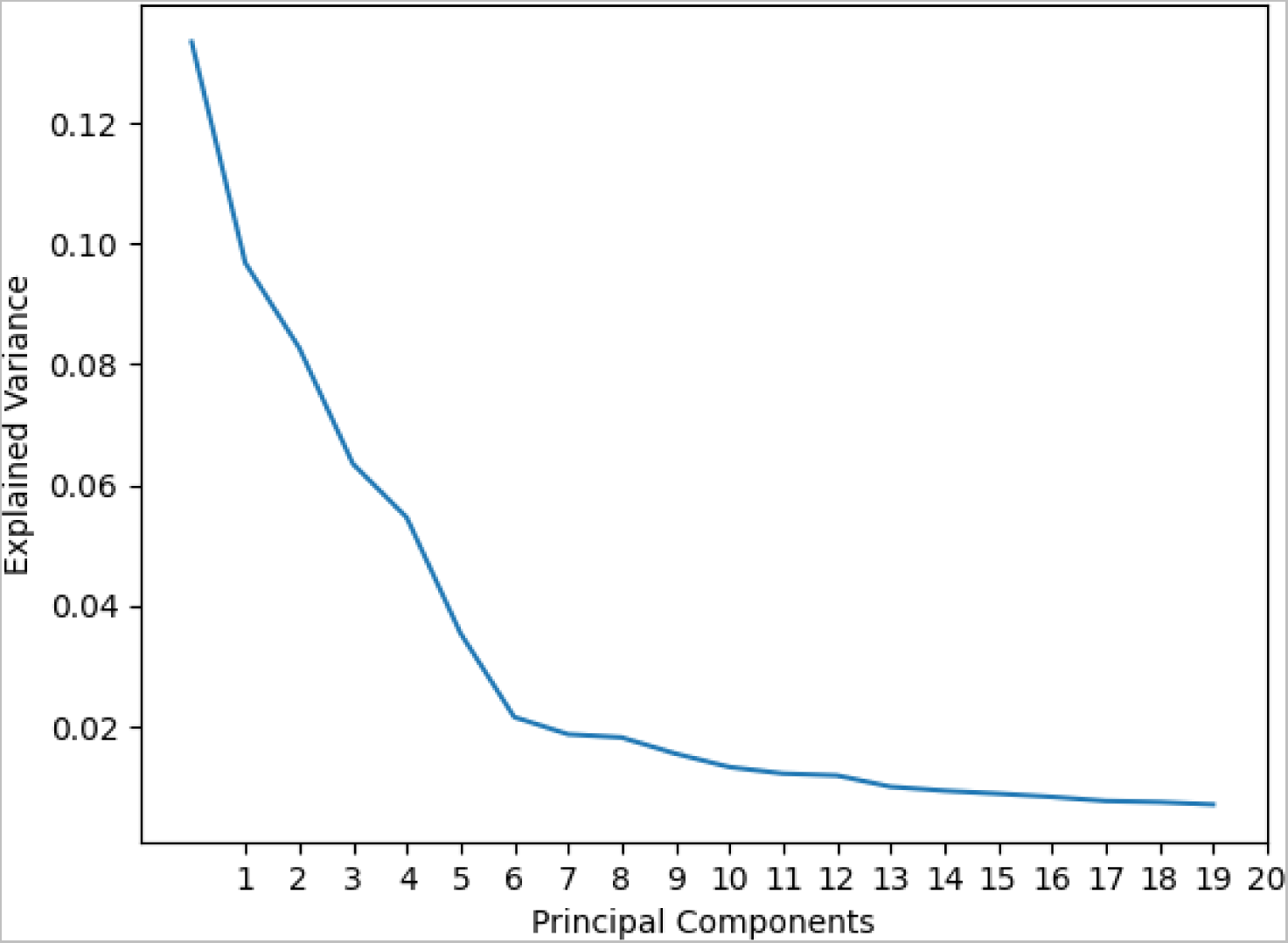
Explainable variances across 20 PCs.

**Supplementary Figure S11.**
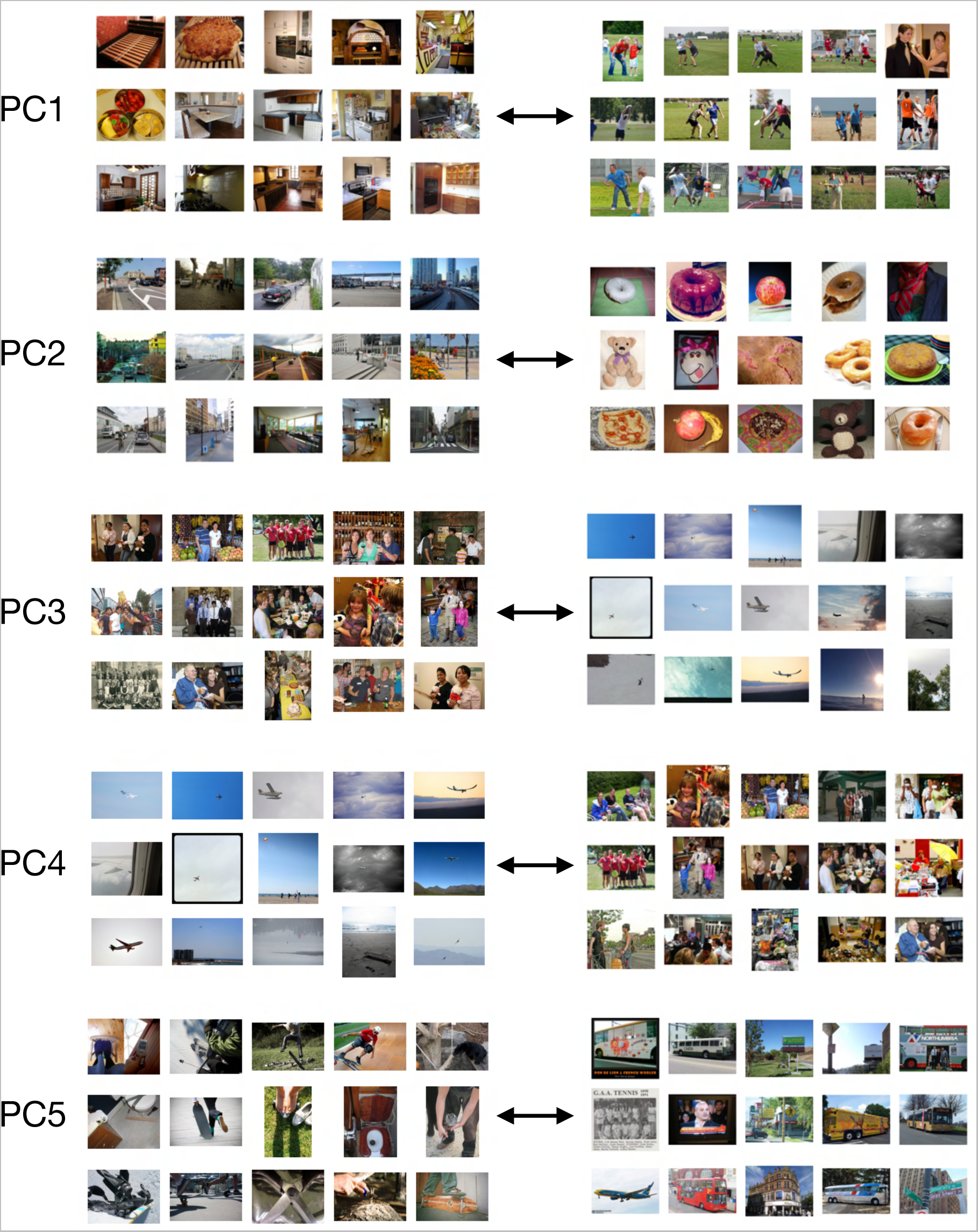
Top 15 images for top 5 PCs. These images were identified by computing the dot product similarity of ResNet*_CLIP_*image embeddings with each PC. Among 10,000 randomly sampled images, the images with most and least similarity within each PC were used to interpret the information encoded in that PC.

**Supplementary Figure S12.**
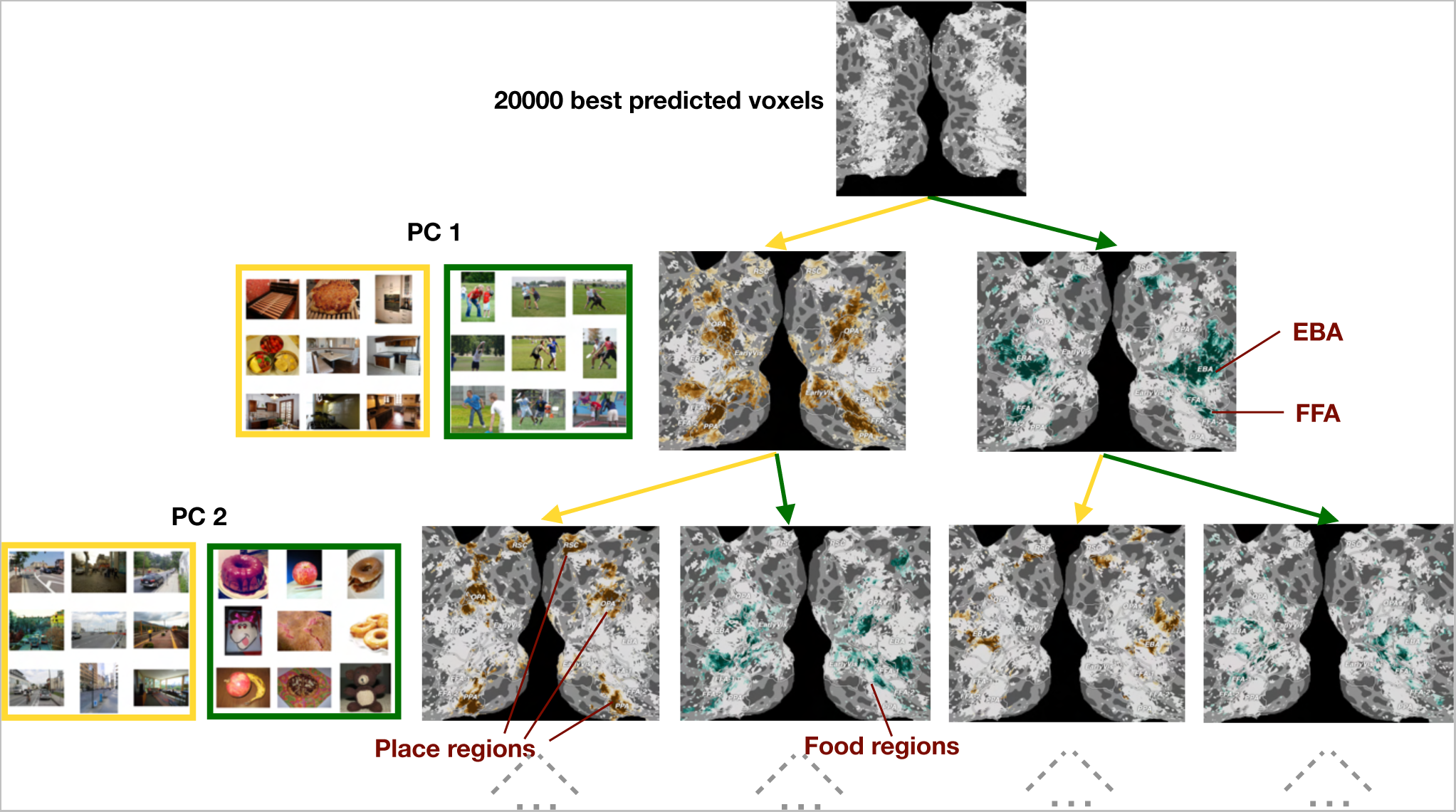
Cortical semantic organization as revealed by the principal components of the CLIP encoding model. Brain regions well predicted by ResNet*_CLIP_* can be hierarchically decomposed using the model PCs. PC1 separates animacy regions (EBA and FFA) from other regions, which are themselves separated by PC2 into place and food regions^40^. The rest of the tree is not shown due to space constraints.

**Supplementary Figure S13.**
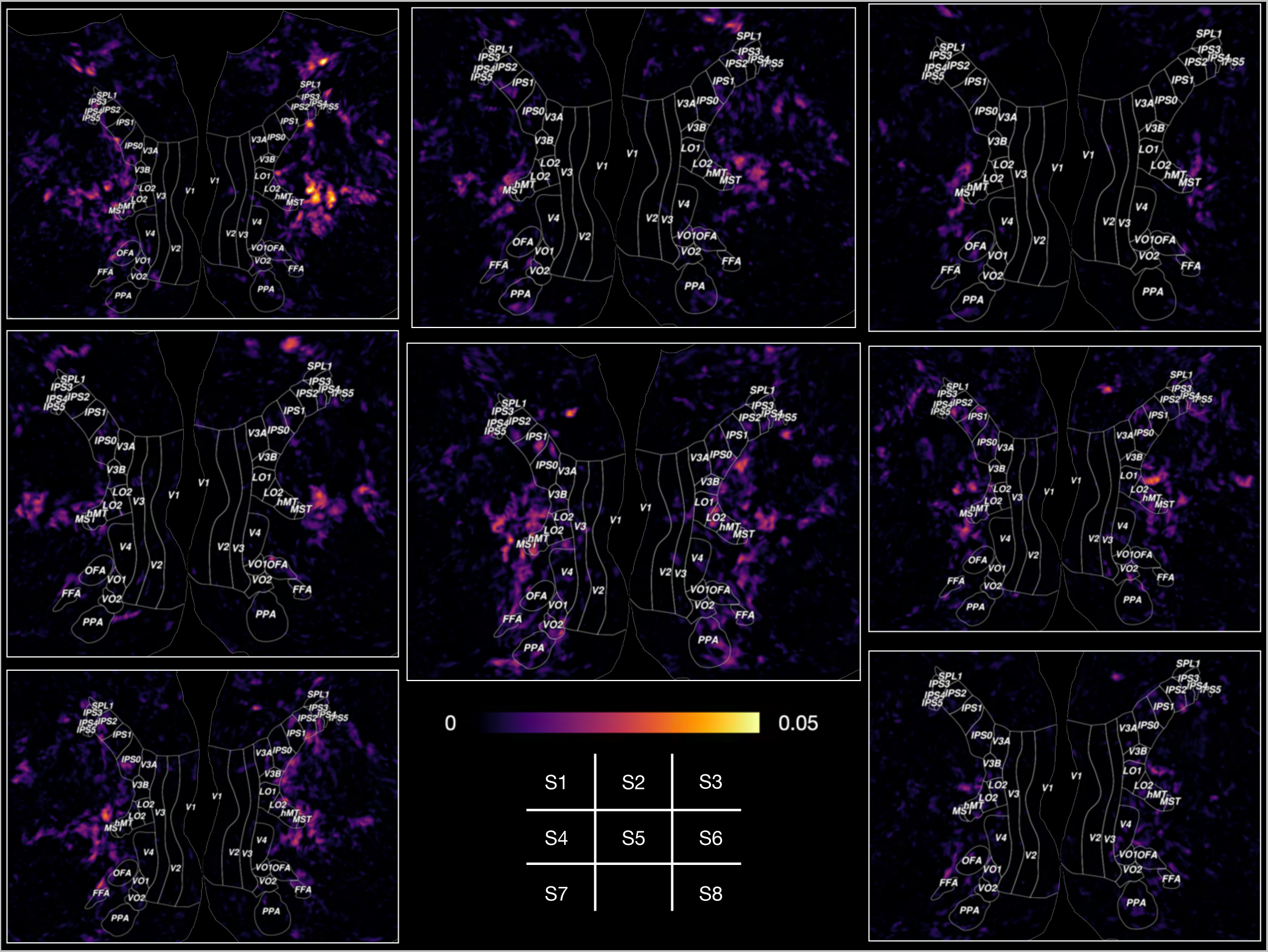
Unique variances by YFCC SLIP compared to YFCC simCLR across all 8 subjects in MNI space. Unique variances explained by YFCC SLIP from each subjects are projected into MNI space. Only unique variances above significance threshold are projected (*p* < 0.05, FDR-corrected^29^.

## References

1. LeCun, Y., Bengio, Y. & Hinton, G. Deep learning. Nature 521, 436–444 (2015).

2. Yamins, D. L. K. et al. Performance-optimized hierarchical models predict neural responses in higher visual cortex. Proc Natl Acad Sci U S A 111, 8619–8624 (2014).

3. Yamins, D. L. K. & DiCarlo, J. J. Using goal-driven deep learning models to understand sensory cortex. Nat Neurosci 19, 356–365 (2016).

4. Toneva, M., Mitchell, T. M. & Wehbe, L. Combining computational controls with natural text reveals aspects of meaning composition. Nat. computational science 2, 745–757 (2022).

5. Deng, J., et al. ImageNet: A large-scale hierarchical image database. In 2009 IEEE conference on computer vision and pattern recognition, 248–255 (IEEE, 2009).

6. Aminoff, E. M. & Tarr, M. J. Associative processing is inherent in scene perception. PLOS ONE 10, 1–19 (2015).

7. Gauthier, I., James, T. W., Curby, K. M. & Tarr, M. J. The influence of conceptual knowledge on visual discrimination. Cogn Neuropsychol 20, 507–523 (2003).

8. Schaffner, J., Bao, S. D., Tobler, P. N., Hare, T. A. & Polania, R. Sensory perception relies on fitness-maximizing codes. *Nat*. Hum. Behav. (2023).

9. Lupyan, G., Thompson-Schill, S. L. & Swingley, D. Conceptual penetration of visual processing. Psychol. Sci. 21, 682–691 (2010).

10. Radford, A., et al. Learning transferable visual models from natural language supervision. In International Conference on Machine Learning (ICML), 8748–8763 (PMLR, 2021).

11. Li, L. H. et al. Grounded language-image pre-training. In 2022 IEEE/CVF Conference on Computer Vision and Pattern Recognition (CVPR), 10955–10965 (2022).

12. Yuan, L., et al. Florence: A new foundation model for computer vision, DOI: 10.48550/ARXIV.2111.11432 (2021).

13. Jia, C., et al. Scaling up visual and vision-language representation learning with noisy text supervision. In International Conference on Machine Learning (ICML) (PMLR, 2021).

14. Wu dao 2.0. https://gpt3demo.com/apps/wu-dao-20. Accessed: 2022-10-20.

15. Pinker, S. *The language instinct: How the mind creates language* (HarperCollins, New York, NY, 2007).

16. Fang, A., et al. Data determines distributional robustness in contrastive language image pre-training (CLIP). In Chaudhuri, K., et al. (eds.) International Conference on Machine Learning, ICML 2022, 17-23 July 2022, Baltimore, Maryland, USA, vol. 162 of Proceedings of Machine Learning Research, 6216–6234 (PMLR, 2022).

17. Mu, N., Kirillov, A., Wagner, D. & Xie, S. SLIP: Self-supervision meets language-image pre-training. In Computer Vision – ECCV 2022: 17th European Conference, Tel Aviv, Israel, October 23–27, 2022, Proceedings, Part XXVI, 529–544 (Springer-Verlag, Berlin, Heidelberg, 2022).

18. Li, L. H., Yatskar, M., Yin, D., Hsieh, C.-J. & Chang, K.-W. VisualBERT: A simple and performant baseline for vision and language, DOI: 10.48550/arXiv.1908.03557 (2019).

19. Tan, H. & Bansal, M. LXMERT: Learning cross-modality encoder representations from transformers. In Inui, K., Jiang, J., Ng, V. & Wan, X. (eds.) EMNLP/IJCNLP (1), 5099–5110 (Association for Computational Linguistics, 2019).

20. Murray, S. O., Boyaci, H. & Kersten, D. The representation of perceived angular size in human primary visual cortex. Nat Neurosci 9, 429–434 (2006).

21. Gilbert, C. D. & Li, W. Top-down influences on visual processing. Nat. Rev. Neurosci. 14, 350–363 (2013).

22. He, K., Zhang, X., Ren, S. & Sun, J. Deep residual learning for image recognition. In Proceedings of the IEEE conference on computer vision and pattern recognition, 770–778 (2016).

23. Devlin, J., Chang, M., Lee, K. & Toutanova, K. BERT: pre-training of deep bidirectional transformers for language understanding. In Burstein, J., Doran, C. & Solorio, T. (eds.) Proceedings of the 2019 Conference of the North American Chapter of the Association for Computational Linguistics: Human Language Technologies, NAACL-HLT 2019, Minneapolis, MN, USA, June 2-7, 2019, Volume 1 (Long and Short Papers), 4171–4186 (Association for Computational Linguistics, 2019).

24. Naselaris, T., Kay, K. N., Nishimoto, S. & Gallant, J. L. Encoding and decoding in fMRI. Neuroimage 56, 400–410 (2011).

25. Allen, E. J. et al. A massive 7T fMRI dataset to bridge cognitive neuroscience and artificial intelligence. Nat. Neurosci. 25, 116–126 (2022).

26. Chen, T., Kornblith, S., Norouzi, M. & Hinton, G. A simple framework for contrastive learning of visual representations. In International Conference on Machine Learning (ICML) (JMLR.org, 2020).

27. Schuhmann, C. et al. LAION-5B: An open large-scale dataset for training next generation image-text models. In Koyejo, S., et al. (eds.) Advances in Neural Information Processing Systems, vol. 35, 25278–25294 (Curran Associates, Inc., 2022).

28. Thomee, B. et al. YFCC100M: The new data in multimedia research. Commun. ACM 59, 64–73 (2016).

29. Benjamini, Y. & Hochberg, Y. Controlling the false discovery rate: a practical and powerful approach to multiple testing. J. Royal statistical society: series B (Methodological) 57, 289–300 (1995).

30. Güçlü, U. & van Gerven, M. A. Deep neural networks reveal a gradient in the complexity of neural representations across the ventral stream. J. Neurosci. 35, 10005–10014 (2015).

31. Huth, A. G., De Heer, W. A., Griffiths, T. L., Theunissen, F. E. & Gallant, J. L. Natural speech reveals the semantic maps that tile human cerebral cortex. Nature 532, 453–458 (2016).

32. Epstein, R. A. & Baker, C. I. Scene perception in the human brain. Annu. Rev. Vis. Sci. 5, 373–397 (2019).

33. Downing, P. E., Jiang, Y., Shuman, M. & Kanwisher, N. A cortical area selective for visual processing of the human body. Science 293, 2470–2473 (2001).

34. Sergent, J., Ohta, S. & MacDonald, B. Functional neuroanatomy of face and object processing: A positron emission tomography study. Brain 115, 15–36 (1992).

35. Kanwisher, N., McDermott, J. & Chun, M. M. The fusiform face area: a module in human extrastriate cortex specialized for face perception. J Neurosci 17, 4302–4311 (1997).

36. Lescroart, M. D., Stansbury, D. E. & Gallant, J. L. Fourier power, subjective distance, and object categories all provide plausible models of bold responses in scene-selective visual areas. Front. computational neuroscience 9, 135 (2015).

37. de Heer, W. A., Huth, A. G., Griffiths, T. L., Gallant, J. L. & Theunissen, F. E. The hierarchical cortical organization of human speech processing. J. Neurosci. 37, 6539–6557 (2017).

38. Saxe, R. & Kanwisher, N. People thinking about thinking people: the role of the temporo-parietal junction in “theory of mind”. In Social neuroscience, 171–182 (Psychology Press, 2013).

39. Çukur, T., Nishimoto, S., Huth, A. G. & Gallant, J. L. Attention during natural vision warps semantic representation across the human brain. *Nat*. neuroscience 16, 763–770 (2013).

40. Jain, N. et al. Selectivity for food in human ventral visual cortex. Commun. Biol. 6, 175 (2023).

41. Pennock, I. M. L. et al. Color-biased regions in the ventral visual pathway are food selective. Curr. Biol. 33, 134–146.e4 (2023).

42. Khosla, M., Apurva Ratan Murty, N. & Kanwisher, N. A highly selective response to food in human visual cortex revealed by hypothesis-free voxel decomposition. Curr. Biol. 32, 1–13 (2022).

43. Conwell, C., Prince, J. S., Hamblin, C. J. & Alvarez, G. A. Controlled assessment of clip-style language-aligned vision models in prediction of brain & behavioral data. In ICLR 2023 Workshop on Mathematical and Empirical Understanding of Foundation Models.

44. Conwell, C., Prince, J. S., Alvarez, G. A. & Konkle, T. Large-scale benchmarking of diverse artificial vision models in prediction of 7T human neuroimaging data. bioRxiv (2022).

45. Conwell, C., Prince, J., Alvarez, G., Konkle, T. & Kay, K. Opportunistic experiments on a large-scale survey of diverse artificial vision models in prediction of 7t human fMRI data. In Conference on Cognitive Computational Neuroscience (2022).

46. Bracci, S. & Op de Beeck, H. P. Understanding human object vision: A picture is worth a thousand representations. Annu. Rev. Psychol. 74, 113–135 (2023).

47. Maier, M. & Abdel Rahman, R. No matter how: Top-down effects of verbal and semantic category knowledge on early visual perception. Cogn. Affect. & Behav. Neurosci. 19, 859–876 (2019).

48. Charest, I., Allen, E., Wu, Y., Naselaris, T. & Kay, K. Precise identification of semantic representations in the human brain. J. Vis. 20, 539–539 (2020).

49. Devereux, B. J., Clarke, A. & Tyler, L. K. Integrated deep visual and semantic attractor neural networks predict fMRI pattern-information along the ventral object processing pathway. Sci. Rep. 8, 10636 (2018).

50. Nappa, R., Wessel, A., McEldoon, K. L., Gleitman, L. R. & Trueswell, J. C. Use of Speaker’s Gaze and Syntax in Verb Learning. Lang. Learn. Dev. 5, 203–234 (2009).

51. Waxman, S. R. & Markow, D. B. Words as invitations to form categories: evidence from 12- to 13-month-old infants. Cogn. Psychol. 29, 257–302 (1995).

52. Lupyan, G., Rakison, D. H. & McClelland, J. L. Language is not just for talking: redundant labels facilitate learning of novel categories. Psychol. Sci. 18, 1077–1083 (2007).

53. Shusterman, A. & Spelke, E. Language and the Development of Spatial Reasoning. In Carruthers, P., Laurence, S. & Stich, S. (eds.) The Innate Mind: Structure and Contents, 89–106 (Oxford University Press, 2005).

54. Lin, T. Y. et al. Microsoft COCO: Common objects in context. Lect. Notes Comput. Sci. (including subseries Lect. Notes Artif. Intell. Lect. Notes Bioinformatics) 8693 LNCS, 740–755 (2014).

55. Dale, A. M., Fischl, B. & Sereno, M. I. Cortical surface-based analysis: I. segmentation and surface reconstruction. NeuroImage 9, 179–194 (1999).

56. Fischl, B., Sereno, M. I. & Dale, A. M. Cortical surface-based analysis: II: Inflation, flattening, and a surface-based coordinate system. NeuroImage 9, 195–207 (1999).

57. Gao, J. S., Huth, A. G., Lescroart, M. D. & Gallant, J. L. Pycortex: an interactive surface visualizer for fMRI. *Front*. Neuroinformatics 9 (2015).

58. Koushik, J. torch-gel. https://github.com/jayanthkoushik/torch-gel (2017).

